# From cooperation to predation: the evolutionary origins of phagocytosis

**DOI:** 10.64898/2026.10.07.757284

**Authors:** Josephine Solowiej-Wedderburn, Alfred Leimar, Christopher Kempes, Jordan G. Okie, Eric Libby

## Abstract

A major transition in the evolution of eukaryotic cells was phagocytosis: the ability of one cell to engulf another. In some cases, the engulfed cell persists, forming an endosymbiosis which is often mutually beneficial. However, it is unclear whether the interaction between the ancestors was also cooperative. There are clear benefits to developing the ability to eat a competitor, but can it ever be beneficial to eat a cooperator? We use a dynamical systems framework to identify the ecological conditions under which phagocytosis can evolve from a system of cooperating microbes. Beginning with a system of two cooperating species, we use invasion analysis to identify the parameter regimes under which a phagocytosing mutant can invade and the subsequent population dynamics. We identify a critical threshold that determines whether an invading predator will coexist with the cooperative community or replace members. Repeatedly invading with different predators, we find that the system saturates when there are three coexisting species, though this community does not have a single optimal composition. However, if we incorporate additional costs, we find there is a single evolutionarily stable community of two cooperators and a pure predator. Ultimately, we show that it is possible for phagocytosis to evolve from a cooperative community and, moreover, coexist with the original cooperating species.

**Highlights:**

- Predation can evolve from a cooperative ecology, as long as the prey species is sufficiently abundant.
- A predatory species can coexist with its ancestral cooperative community.
- There is a niche separation with one species displaying more predatory and the other more cooperative behaviour.
- Imposing a cost to mixed strategies leads to a unique ecologically and evolutionarily stable community consisting of specialist species (the original two cooperators and a pure predator).

## 1. Introduction

The origin of eukaryotes is a major transition in evolution that facilitated the rise of a diverse array of complex macroscopic life (Szathmáry and Smith, 1995). A pivotal event during the evolution of eukaryotes was an endosymbiosis between two prokaryotes, an endosymbiotic Alphaproteobacterium that evolved into mitochondria and an archaean that hosted the endosymbiont (Vosseberg et al., 2024). While endosymbiosis is common in eukaryotes, it is exceptionally rare in extant prokaryotic hosts. One possible reason for the rarity is the observation that very few known prokaryotes possess a mechanism to engulf another cell, say through phagocytosis, thereby preventing initial establishment of an endosymbiosis (Lage et al., 2025). This has led some to suggest that the evolution of phagocytosis was a precursor to endosymbiosis in the evolution of eukaryotes (Cavalier-Smith, 2002; Martijn and Ettema, 2013; Zachar et al., 2018). Yet, at some point it evolved, which begs the question: what conditions facilitated the evolution of phagocytosis?

Both intrinsic and extrinsic factors could limit the evolution of phagocytosis. A key intrinsic factor is that certain cytoskeletal architectures may not permit a cell to engulf another. Indeed, eukaryotes and prokaryotes differ significantly in their interior cell structure and organization which suggests that these changes are deep-seated and may have required many evolutionary steps. Thus, as the ancestors of eukaryotes were evolving a cytoskeletal architecture that diverged from prokaryotes, there must have been many factors instead of phagocytosis driving this process. Even once cells possessed an architecture that could allow for phagocytosis, extrinsic factors such as environmental or ecological conditions would have played a role in determining whether or not phagocytosis would evolve. For example, consider the analogous case of genetic competence in prokaryotes. Many prokaryotes have mechanisms for importing DNA from their external environment but these mechanisms are regulated to occur only in specific circumstances. Similarly while a cytoskeletal architecture may permit phagocytosis there may be particular extrinsic conditions needed to actively select for the evolution of phagocytosis.

The question of what conditions favour the evolution of phagocytosis is paradoxical when framed in the context of the evolution of eukaryotes via endosymbiosis. In order for phagocytosis to be advantageous one cell must benefit from engulfing another and digesting it. Thus, there is the potential for two types of benefit: 1. removing another cell from the population and 2. gaining resources from digesting that cell. Both benefits can occur if the cells are in an environment where they compete for the same resources (Ispolatov et al., 2023). If phagocytosis always resulted in the complete digestion of a cell, there would never be the chance to establish an endosymbiosis. Yet, if for some reason a cell engulfed another but did not digest it, the endosymbiont would be a competitor whose survival would be unlikely to afford a fitness benefit to the host (for a possible exception see (Zachar et al., 2018)). This has led some to suggest that phagocytosis evolved late during eukaryogenesis, and after the future host and endosymbiont cells had already evolved a mutually beneficial relationship, e.g., a syntrophy (Baum and Baum, 2014; Imachi et al., 2020; Mills, 2020). However, if two species have a mutually beneficial relationship then it seems unlikely for one to evolve to eat the other.

Here, we use mathematical models to determine whether phagocytosis can evolve from cooperation. We model the cooperation between two species in which one species produces a self-inhibitory by-product that is also a resource for a second species, ensuring that they always grow better together than alone, inspired by syntrophies typically observed in anoxic environments such as deep-sea vents (Morris et al., 2013). We use consumer-resource models to represent these two populations and invasion analysis to determine the conditions for a phagocytosing mutant to invade. To represent the evolution of phagocytosis from the self-inhibitory ancestor, we introduce mixotrophic mutants that predate with predation fraction, *p*, and behave like their ancestor with the remainder, (1−*p*). If *p* = 1, then the mutant is a pure predator. We find that the success of an invasion is independent of *p*, but there is a critical threshold that determines whether the mutant will coexist with or replace its ancestor. We further assess the evolutionary response of the prey species and subsequent invasion of phagocytosing mutants. In all cases, we find that the system saturates when there are three coexisting species leading to evolutionary neutrality, and a resulting community that does not have a single optimal composition. Incorporating an additional cost to the switching behaviour of mutants eliminates this neutrality and produces a single evolutionarily-stable community with the original two cooperators and a pure predator. Thus, our work indicates that it is possible for phagocytosis to evolve from and coexist with a cooperative community.

## 2. Model

### Cooperation by the consumption of an inhibitory by-product

To determine whether a predatory behaviour can evolve from a cooperative interaction, we begin with a simple system of two species, *X* and *Y* . Each species has their own growth resources, *R*_*X*_ and *R*_*Y*_ . Motivated by the wide diversity of microbial syntrophies found in anoxic environments (Morris et al., 2013), such as deep-sea vents, and the various observed and theoretically proposed syntrophies and other cooperative relationships between Alphaproteobacteria and archaea (Spang et al., 2019; López-García and Moreira, 2019, 2023; Mills et al., 2022), we implement cooperation via the production of an inhibitory by-product. When species *X* consumes its resource *R*_*X*_, it produces a waste product *B*. This waste product binds to *R*_*X*_ <u>f</u>orming an inedible compound 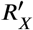 thereby reducing resource availability for *X*. Species *Y* can use this waste product *B* as an alternative resource for growth, establishing a cooperative interaction between the two species. We assume the system is in a well-mixed chemostat with constant outflux, as illustrated in figure 1, and so can be modelled by the following series of differential equations:

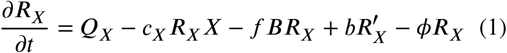

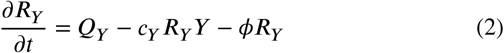

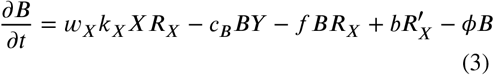

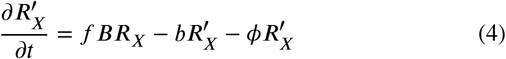

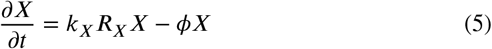

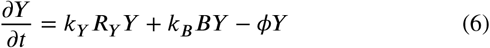

where we assume that all parameters, described in table 1, are ≥ 0. We additionally rescale time by the outflux rate *ϕ* introducing new variable *τ*:= *ϕt/*, outlined in SI 1. For simplicity, we use *τ* and time-rescaled parameters in our subsequent analyses.

**Table 1:** Parameter table.

| parameter | time rescaling | interpretation |
| --- | --- | --- |
| $Q_X$ | $/\phi$ | influx of $R_X$ |
| $Q_Y$ | $/\phi$ | influx of $R_Y$ |
| $c_X$ | $/\phi$ | rate at which $X$ feeds on $R_X$ per units biomass |
| $c_Y$ | $/\phi$ | rate at which $Y$ feeds on $R_Y$ per units biomass |
| $c_B$ | $/\phi$ | rate at which $Y$ feeds on $B$ per units biomass |
| $f$ | $/\phi$ | rate at which inhibitory by-product $B$ binds to $R_X$ |
| $b$ | $/\phi$ | rate at which $R'_X$ dissociates back to $R_X$ and $B$ |
| $k_X$ | $/\phi$ | growth rate of $X$ when consuming $R_X$ per units biomass |
| $k_Y$ | $/\phi$ | growth rate of $Y$ when consuming $R_Y$ per units biomass |
| $k_B$ | $/\phi$ | growth rate of $Y$ when consuming $B$ per units biomass |
| $w_X$ | n/a | ratio of mass of $B$ produced relative to $X$ biomass production when consuming $R_X$ |
| $\phi$ | n/a | constant outflux rate |

**Figure 1:**
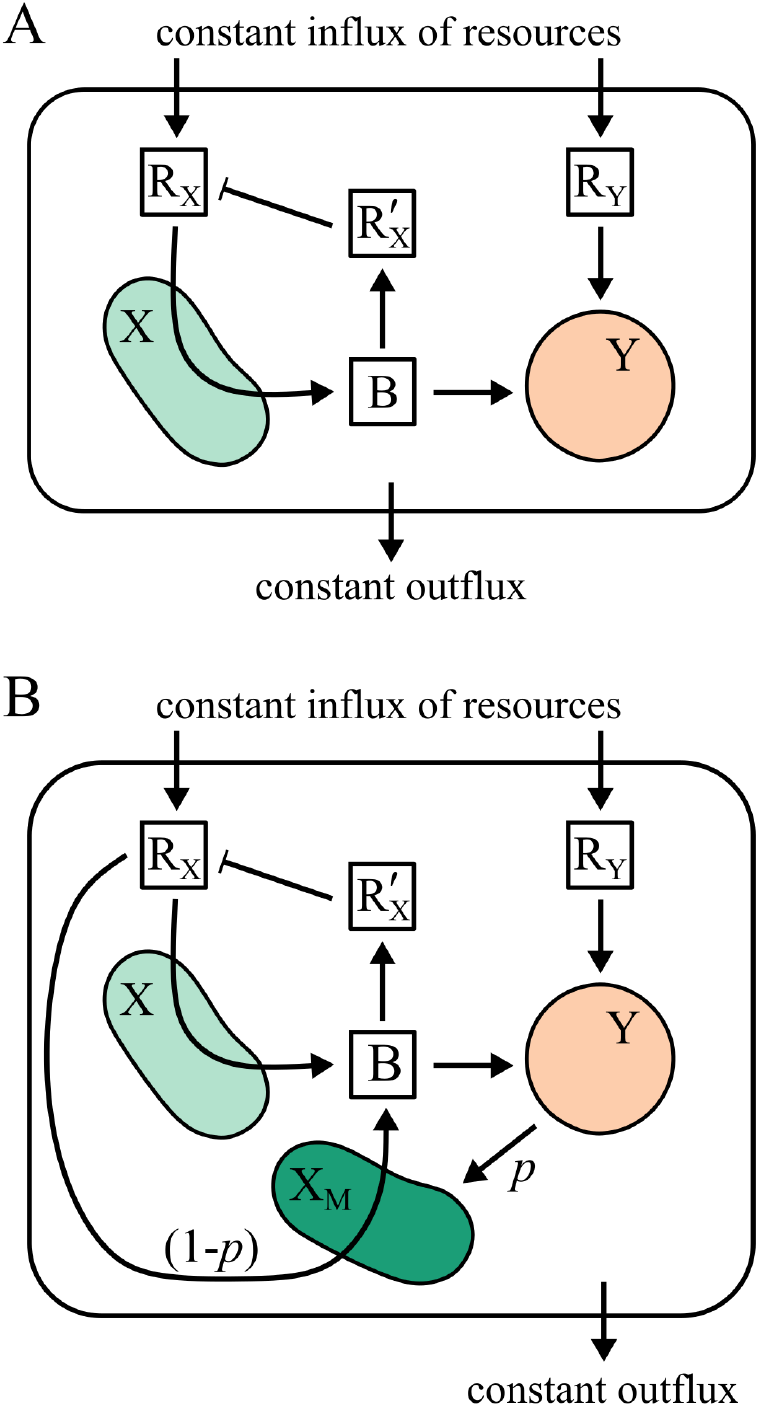
Models for cooperation and phagocytosis. A) Schematic shows cooperative interaction between two species *X a*nd *Y* . Species *X c*onsumes its resource *R*_*X*_, producing a waste product *B t*hat acts as an inhibitor, binding to *R*_*X*_ *t*o form an indigestible state 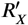. Species *Y c*onsumes its resource *R*_*Y*_, but it can also consume by-product *B*. Thus *Y b*enefits from the presence of *X* as it gains a new resource; and *X* benefits from the presence of *Y* as it removes/reduces the inhibitor *B*. We consider the system in a chemostat with constant resource influxes, constant outflux, and well-mixed environment. B) Schematic shows model for the evolution of phagocytosis from initial cooperative ecology. The interaction between cooperative species *X* and *Y* is the same as in (A), but now there is a new mutant species *X*_*M*_ . Species *X*_*M*_ consumes *Y* via phagocytosis with predation fraction *p* and behaves like its ancestor *X* with remainder (1 − *p*).

In SI 2 we solve the system of ODEs (1)–(6) (rescaled in time) at steady state and find four biologically possible equilibria: the trivial case when neither *X* nor *Y* grow; two cases corresponding to when *X* or *Y* grow in isolation; and one case when *X* and *Y* coexist. Since we are interested in cooperation, we focus on the coexistence state and parameters that cause this state to be stable. Specifically, species *X* can survive if *k*_*x*_*Q*_*x*_ > 1. Species *Y* has two possible resources, *R*_*Y*_ and *B*, each of which implies a different condition for *Y* to survive. If *k*_*y*_*Q*_*y*_ > 1, then there is sufficient resource *R*_*Y*_ for *Y* to survive even in the absence of *B*. For *Y* to survive solely on *B*, there is a further condition (see SI 2) on *k*_*x*_*Q*_*x*_ to ensure that there is both enough resource for *X* to survive on *R*_*X*_ and produce sufficient waste product for *Y* .

In regimes where *k*_*y*_*Q*_*y*_ < 1, but *k*_*x*_*Q*_*x*_ is sufficient to sustain both *X* and *Y* populations, *Y* is obligate on *X*. If *X* were to be obligate on *Y*, it would need to produce so much by-product *B* that it would block its uptake of *R*_*X*_. However, since *B* is produced after uptake of *R*_*X*_, this is not possible. This can be seen in the condition for *X* to survive, *k*_*x*_*Q*_*x*_ > 1, which does not have any dependence on *B*. In SI 3 we show how the dynamical system (1)–(6) can be adapted to include an external influx of inhibitory compound *B*. We show that in this more general system there are parameter regimes in which *X* is obligate on *Y* . Biologically, this could arise if, for example, *X* and *Y* are part of a larger microbial community where other members also produce by-product *B*. As we focus on the interaction between *X* and *Y*, we restrict our analyses to the original system with zero influx of *B*.

## 3. Results

### Successful invasion of a predatory mutant

For these analyses we use the time-rescaled system of ODEs (S1)–(S6) that describe cooperation between two species, *X* and *Y* . In this system *X* produces a self-inhibitory by-product *B* which can be consumed by species *Y* with a growth benefit. Since we are interested in whether phagocytosis can evolve from a cooperative interaction, we add a mutant species *X*_*M*_ that predates on *Y* to the system. In the equations this predatory interaction is represented by a product term similar to those used in Lotka-Volterra equations. This captures the internal digestion of prey, as opposed to external digestion which might generate public goods. We model *X*_*M*_ as a mixotroph, engaging in predation with predation fraction *p* and behaving like *X* with remainder (1 −*p*). In the extreme case *p* = 1, *X*_*M*_ is a pure predator feeding only on *Y* . When *X*_*M*_ behaves like *X*, we assume that it has exactly the same parameters for consumption of *R*_*X*_, production of *B*, and growth. This introduces additional terms to the equations for the dynamics of *R*_*X*_ and *B*, while predation on *Y* also changes the dynamics of the *Y* population. The resulting system is:

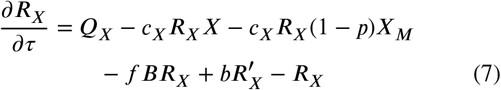

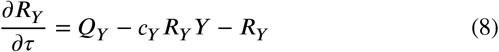

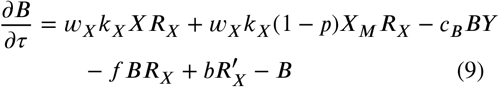

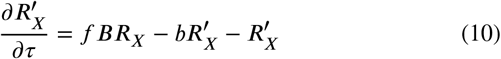

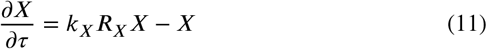

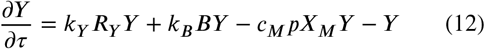

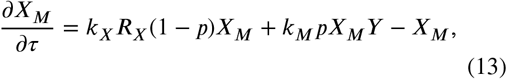

where we have new parameters *c*_*M*_ and *k*_*M*_ that quantify the rate *X*_*M*_ feeds on *Y* and its subsequent growth benefit. We note that if the mutant population *X*_*M*_ is 0, then we get back to the original ODE system for two cooperators, *X* and *Y* .

We use an adaptive dynamics (Brännström et al., 2013) approach to assess whether the mutant species can invade the two-cooperator system. In adaptive dynamics, it is assumed that a small population of mutants *X*_*M*_ enter the system at steady state and their initial population dynamics are exponential, *X*_*M*_ = *eλτ* . Hence, the initial growth rate is given by

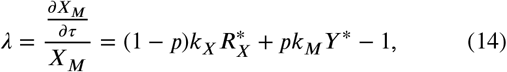

where ^*^-ed variables represent the two-cooperator steady state solutions. We evaluated these steady states and found 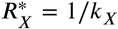 when *X* and *Y* coexist (see SI 2). Substituting this into expression (14) we see that

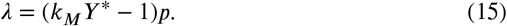

If *λ* > 0 then we expect the initial mutant population will grow exponentially and so invade, while if *λ* < 0 then it cannot invade.

Expression (15) for *λ* shows that the invasion of a mutant is independent of the feeding probability p ∈ (0, 1]. Instead it relies only on there being a sufficient population *Y*^*^ for the mutant to feed on, scaled by *k*_*M*_ the growth benefit to *X*_*M*_ . While the conditions for successful invasion of a mutant are independent of the predation fraction, *p*, the magnitude of *λ* is proportional to *p*. Hence, we expect that the initial growth rate of the invading mutant population to be greater for higher values of *p*, as illustrated in fig. 2A.

**Figure 2:**
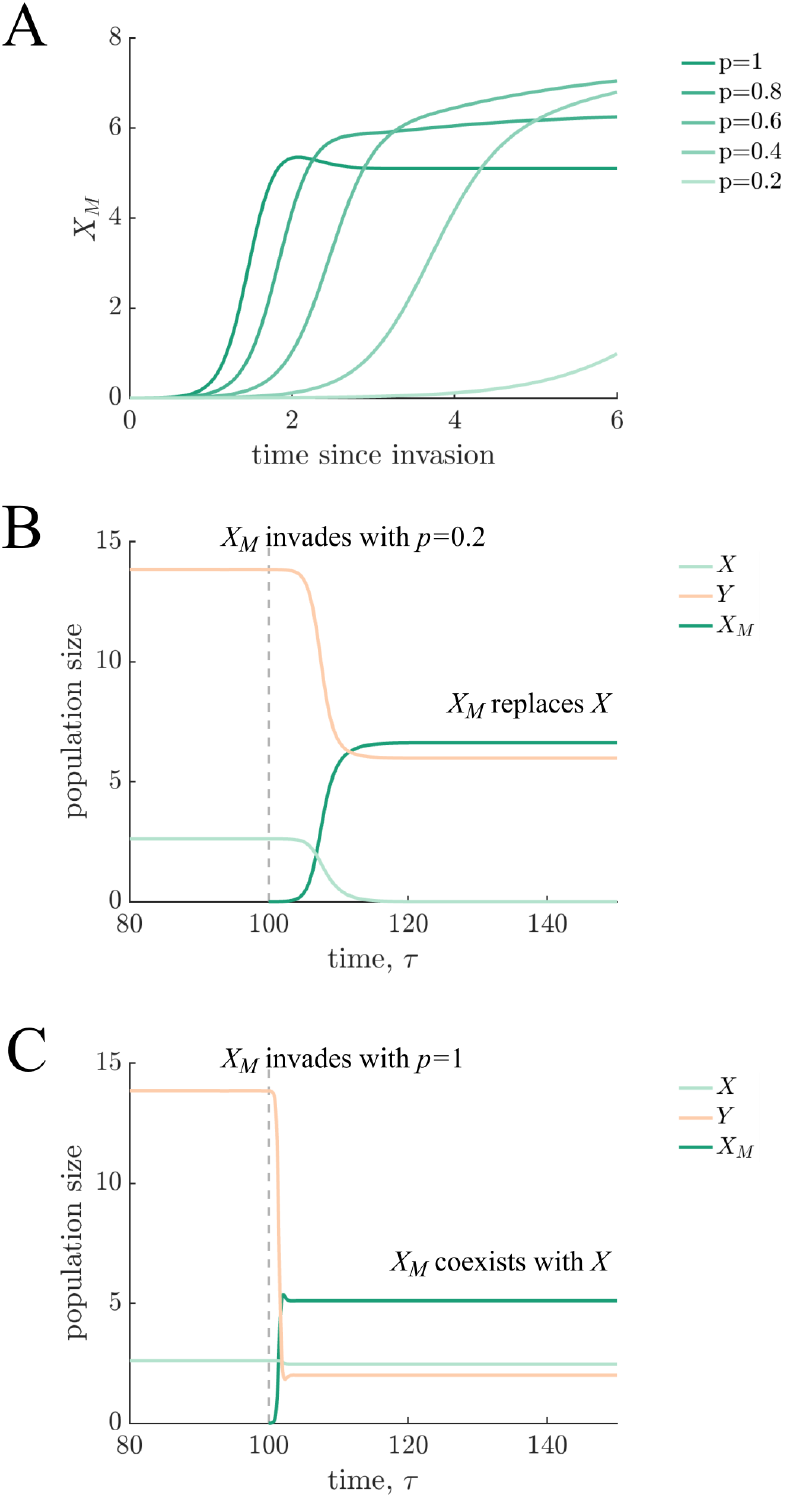
Invasion of a predatory mutant. A) Plot shows the initial dynamics of a mixotrophic mutant *X*_*M*_ with different predation fractions. From lightest to darkest, *p* = 0.2, 0.4, 0.6, 0.8, 1 such that the darkest line represents a pure predator. Other parameters were *Q*_*X*_ = 3.7, *Q*_*Y*_ = 3.2, *c*_*X*_ = 9.0, *c*_*Y*_ = 4.0, *c*_*B*_ = 7.1, *k*_*X*_ = 6.7, *k*_*Y*_ = 1.7, *k*_*B*_ = 3.7, *f* = 5.6, *b* = 3.9, *w*_*X*_ = 9.3 such that the benefit to *X* is ×2 when growing with *Y* and the benefit to *Y* is ×12.5; additional parameters for the mutant are *k*_*M*_ = 0.5 and *c*_*M*_ = 1. The growth rate of the mutant immediately after invasion is higher for higher values of *p*. B and C) Plots show the population dynamics of *X, Y*, and *X*_*M*_ where *X*_*M*_ invades at time *τ* = 100. The parameters are the same as in (A) with two different predation fractions: in (B) *p* = 0.2, and in (C) *p* = 1, a pure predator. In both cases the mutant can invade—this is independent of *p*—however, in (B) *X*_*M*_ replaces the *X* population, while in (C) all three species *X, Y*, and *X*_*M*_ coexist post-invasion.

### Steady states following invasion

In the previous section, we determined the conditions for invasion of a predatory mutant *X*_*M*_ . Here, we consider the consequences of this invasion on the resident species *X* and *Y* . In figs. 2B and C we plot the population dynamics for two examples with exactly the same parameters except different values of *p*. In fig. 2B the invading *X*_*M*_ population replaces *X*, while in fig. 2C all three species coexist post invasion. In SI 4 we identify the steady state solutions with a positive mutant community, *X*_*M*_ > 0. We find that there is a critical threshold *p*_*crit*_ that ensures the coexistence of *X* and *X*_*M*_:

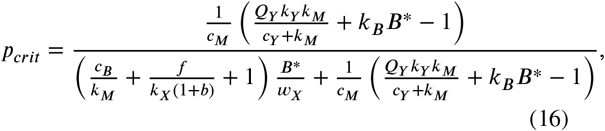

illustrated in fig. 3. If *p* > *p*_*crit*_, then the mutant *X*_*M*_ can coexist with the original cooperative species *X* and *Y* . We note that a pure predator with *p* = 1 can always coexist with *X* and *Y* . However, if *p* ≤ *p*_*crit*_, then the mutant outcompetes the original *X* species for its sole resource *R*_*X*_, driving it extinct.

**Figure 3.**
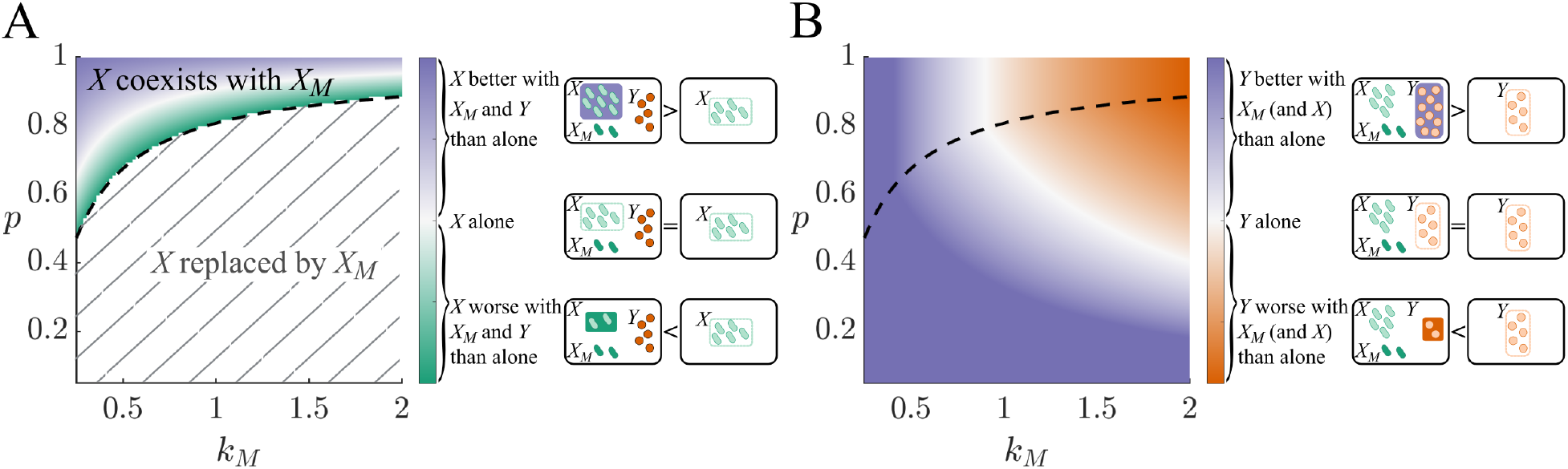
*X* and *Y* populations after invasion. Heat maps show the populations of *X* (A) and *Y* (B) after the successful invasion of a mutant *X*_*M*_ . We vary *p* and *k*_*M*_ and other parameters are as in fig. 2 such that *X*_*M*_ can invade. The dashed line indicates *p*_*crit*_; if *p* f *p*_*crit*_ then *X* is replaced by *X*_*M*_ and if *p* > p_*crit*_ then *X* coexists with *X*_*M*_ and *Y* . The colour bars indicate the relative population sizes of *X* and *Y* in the system with *X*_*M*_ *vs* alone. In both panels purple shading is used to indicate parameter regimes where *X* or *Y* have larger population sizes at steady state despite predation by *X*_*M*_ .

After a successful invasion of mutant *X*_*M*_, both *X* and *Y* populations decrease. The *Y* population is reduced because of predation by *X*_*M*_ . There are two factors that cause the *X* population to be reduced: competition with a mixotrophic *X*_*M*_ and increased concentration of inhibitory by-product *B* owing to a smaller *Y* population. If instead we compare their population sizes with the corresponding single-species systems, we find that there are some parameter regimes where *X* and *Y* can grow better in the three-species system than when grown independently. In fig. 3 we plot heat maps of *X* and *Y* in the three-species steady state normalised by their population sizes in isolation. This shows that when *X*_*M*_ has a high predation fraction and relatively low growth rate on *Y* (i.e. lower *k*_*M*_), it is possible for both *X* and *Y* to achieve higher abundances in the presence of *X*_*M*_ than by themselves.

### Evolutionary response of prey species

Having esablished that a predatory species *X*_*M*_ can invade, we now study the evolutionary response of *Y* . Equation (12) suggests that *Y* could benefit by increasing its growth rate, either from the consumption of its own resource (i.e., increasing *k*_*Y*_) or from the consumption of waste product *B* (i.e., increasing *kB*). As we are considering ecological interactions, we focus on the parameter *k*_*B*_. To relate changes in *k*_*B*_ to mechanistic adaptations in *Y*, we rewrite *k*_*B*_ as 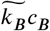, where *c*_*B*_ is the rate of consumption of *B* and 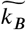 is its conversion to growth. We note that this decomposition will not affect the invasion criteria but it will affect what happens to the other species.

Again, we use adaptive dynamics to evaluate the invasion criteria for a mutant *Y*_*M*_ species with the same growth dynamics as *Y* but new parameter 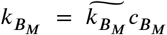 . The initial growth rate of *Y*_*M*_ is given by

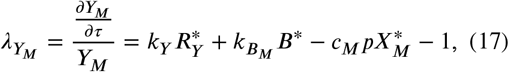

where ^*^ -ed variables represent the steady state solution, assuming a predatory species *X*_*M*_ has already established. We see from expression (17) that if 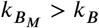 then 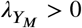. Thus if the mutant *Y*_M_ consumes *B* at a higher rate 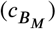 or increases its efficiency in converting resource into growth 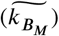, it can invade. When *Y*_*M*_ is able to invade in these circumstances, it replaces *Y* .

The successful invasion of a mutant *Y*_*M*_ has effects on the other species in the community (see fig. 4). For example, if *Y*_*M*_ invades a community with an *X*_*M*_ population, it will cause *X*_*M*_ to reach a higher steady state. In contrast, if *Y*_*M*_ invades a community with an *X* population, it could lead *X* to reach a higher or lower steady state— even go extinct. Which of these outcomes occurs depends on a couple of factors. First, if *X*_*M*_ is not present then the *X* population always benefits from invasion by *Y*_*M*_ . Second, if *X*_*M*_ is present then the consequences of *Y*_*M*_ *’*s invasion on *X* is determined by

**Figure 4:**
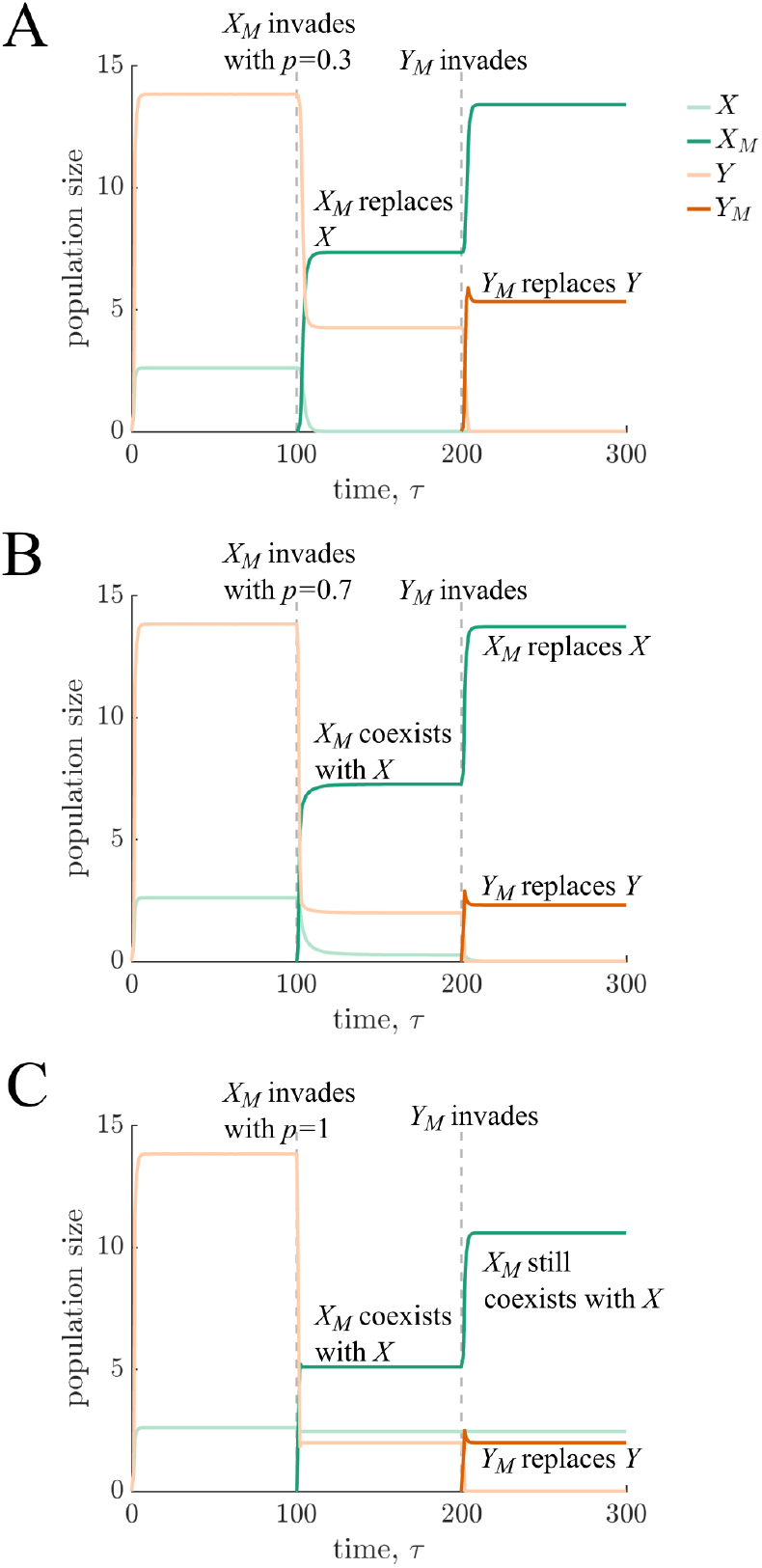
Invasion of *Y*_*M*_ mutant. Plots show the dynamics of the invasion of mutant *Y*_*M*_ with higher cooperative effect *k*_*B,M*_ = 2*k*_*B*_ following the invasion of mutant *X*_*M*_ . Specifically, we illustrate 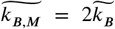, see SI fig. S2 for the effects of adaptation in *c*_*B*_. *X*_*M*_ invades at time *τ* = 100 and *Y*_*M*_ at *τ* = 200. The different panels correspond to *X*_*M*_ mutants with different predation fractions: in (A) *p* = 0.3 < *p*_*crit*_ so *X*_*M*_ replaces *X*, while in (B) *p* = 0.7 > *p*_*crit*_ and (C) *p* = 1 so X_M_ coexists with *X* and *Y* . Other parameters are as in fig. 2. In all cases *Y*_*M*_ replaces *Y* and *X*_*M*_ benefits from the invasion of *Y*_*M*_ . The invasion of the *Y*_*M*_ mutant shifts the critical *p*_*crit*, n*ew*_ driving the *X* population extinct in panel (B).

whether the adaptation in 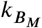 manifests as an adaptation in 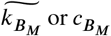 (see analyses in SI 5). Ultimately, these analyses show that a mutant *Y*_M_ that increases its consumption of the waste product *B* can help the mixotrophic species *X*_*M*_*—* the one that is preying on it—more than the original, non-phagocytotic *X* species.

### Invasion of subsequent *X*_*M*_ mutants

We now consider whether further mutant species *X*_*M*_ can invade. As the evolutionary response of *Y* did not change the dimensionality of the system, we no longer consider *Y*_*M*_ and return to the system following the successful invasion of a first mutant 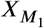 with predation fraction p_1_. We denote 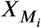 as the i^*th*^ mutant invading the system with predation fraction *p* . We are interested in whether a mutant with a different feeding strategy (*p*_*i*_ *≠ p*_1_) can invade and coexist with or replace any species in the current population. We continue to use an adaptive dynamics approach and consider the growth rate of a new mutant, 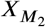, that also follows the dynamics given by differential equation (13), with predation fraction *p*_2_. We assume there is no direct interaction between the 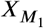 and 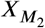 species but they do interact indirectly through consumption of the resource and the *Y* species. Hence, the initial growth rate for this second mutant is also given by expression (14), in terms of the steady state values of resource 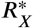 *a*nd species *Y*^*^ before invasion. We note that these steady state values depend on the predation fraction, *p*_1_, adopted by 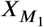 and whether *X* is present.

In the case that the community has all three species, a new invading mutant would enter the system with zero growth rate. This occurs because in order for a community to have three species the first mutant must have had a predation fraction *p*_1_ > *p*_*crit*_. The steady state values for variables *R*_*X*_ and *Y* would then be 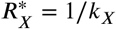 and *Y*^*^ = 1*/k*_*M*_ (see SI 4), which when substituted into expression (14), gives *λ*_2_ = 0. Thus the mutant would not die out, but nor would it be able to grow. The same applies to any further mutants 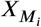 trying to invade with feeding strategy *p*_*i*_ ∈ (0, 1]. Fig. 5B illustrates an example, where all mutant populations 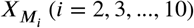 remain at their small initial invasion population values (and are too small to see). The ecological niche is saturated after the invasion of 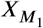 leading to evolutionary neutrality (Diekmann et al., 2002; Geritz et al., 1998).

**Figure 5:**
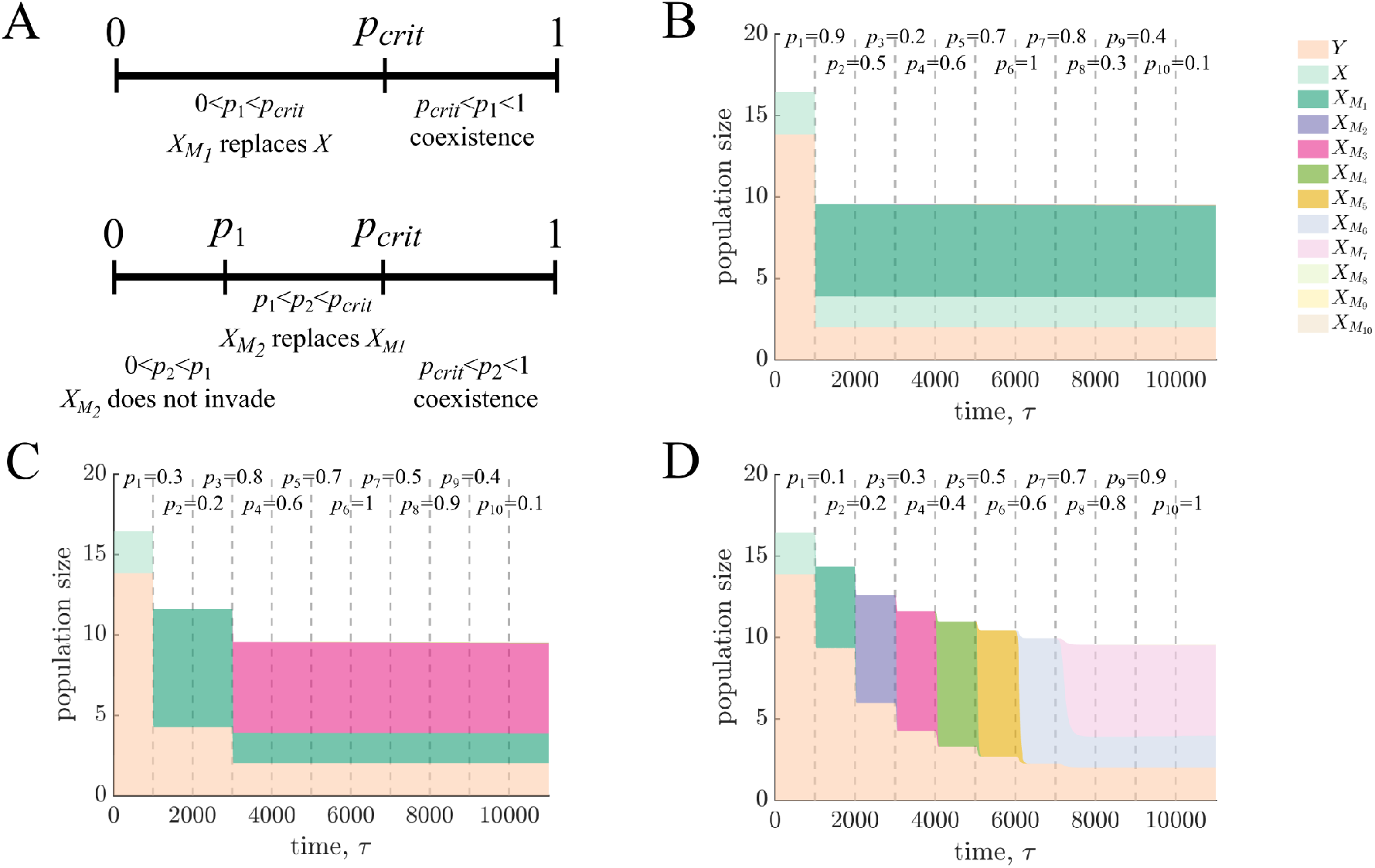
Multiple mutant invasions. A) Schematic summarises the effect of feeding strategy, *p*, on the invasion of mutant species and subsequent community evolution. The successful invasion of the first species is independent of its predation probability, *p*_1_, however, the value of *p*_1_ determines whether the invading 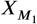 coexists with the existing *X* species or drives it extinct. A second mutant 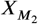 can only invade if *p*_1_ < *p*_crit_ and *p*_2_ > *p*_1_. Again, the community evolution depends on the value of *p*_2_ in relation to *p*_*crit*_ with coexistence occurring when *p*_2_ > *p*_*crit*_. B–D) Plots show examples of simulations with 10 attempted mutation invasion events. Each mutant 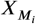 has a different predation fraction, *p*_*i*_ ∈ {0.1, 0.2, …, 1}; all other parameters are as in fig. 2 to ensure mutants can invade and *p*_*crit*_ ≈0.68. A new invasion is attempted after 1,000 time steps to ensure that the existing populations are at steady state. Both the population dynamics following invasion and the success of subsequent invasions depend on the feeding strategy of the invading mutant and existing species. Three different scenarios are shown. In (B) the first mutant to invade has *p*_1_ > *p*_*crit*_ so that it invades and coexists with *X* and *Y* . In (C) *p*_1_ < *p*_*crit*_ so 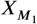 replaces the original *X* population; *p*_2_ < p_1_ so 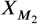 fails to invade; but *p*_*3*_ > *p*_*crit*_ > *p*_1_ so 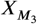 invades and coexists with the existing 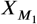 and *Y* populations. In (D) we have subsequent invasions with replacement until *p*_7_ > *p*_*crit*_ which coexists with the existing 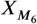 and *Y* populations. In all the examples, once a system with three coexisting species establishes the ecological niche is saturated and results in an evolutionarily stable community.

In the case that the community only has two species, *Y* and 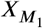, the initial growth rate of a second mutant is

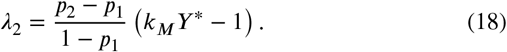

One way for *λ*_2_ to be positive is if *Y*^*^ > 1*/k*_*M*_ and *p*_2_ > *p*_1_. Simulations confirm that *Y*^*^ > 1*/k*_*M*_ when 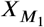 replaces *X* (see fig. 3B). Thus, the second mutant can only invade if it has a higher predation fraction than the first mutant (illustrated in fig. 5C and D). As when 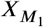 invaded, whether or not a second mutant will coexist with 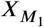 depends on the value of its feeding strategy. In SI 6 we solve the system of ODEs for *Y* with two mutant species 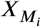 and 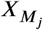 and show that if *pj* > *pcrit* both 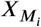 and 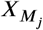 can coexist (illustrated when 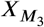 invades in fig. 5C and when 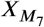 invades in fig. 5D). Meanwhile if *p*_*i*_ < *p*_*j*_ < *p*_*crit*_, then 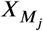 will replace 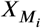(illustrated when 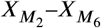 invade in fig. 5D). Fig. 3A summarizes the different ranges of values of *p*_1_ and *p*_2_ for the invasion of two mutants and their implications for the community steady states. Once three species have established a community, the steady state values create another case of evolutionary neutrality, so that *λ*_*i*_ = 0 for all future invasive mutants.

### Mixotrophic switching cost

Our previous analyses showed that when three species coexist the ecological niche was saturated leading to evolutionary neutrality. We also showed that the feeding strategies of the coexisting 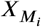 and 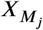 species is not unique. The model that led to these results assumed that mixotrophs can freely switch between resource *R*_*X*_ and predation, allowing for an infinite superposition of mixotrophs with different predation fractions, *p*_*i*_ as long as the total 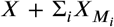 population remains constant (see SI 7). However, feeding on different resources often requires different adaptations and digestive machinery in a cell, hence it might be more biologically realistic to impose a cost to species with a dual feeding strategy. We implement this by modifying the growth rates of mixotrophs by a cost, *s*. This changes the dynamics of the mutant, which we now denote *X*_*S*_ to distinguish the switching cost-imposed mutants from the *X*_*M*_ mutants in earlier analyses. In this model formulation, we also distinguish populations with pure strategies that do not pay a switching cost: *X* that only consumes the resource *R*_*X*_ and *X*_*P*_ that only consumes *Y* . The ODEs for *X*_*S*_ and *X*_*P*_ are given by

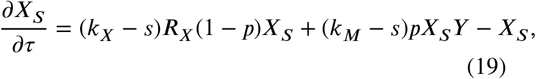

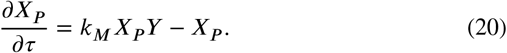

See SI 8 for the full system of equations in this model reformulation.

Invasion of a mutant *X*_*S*_ with a switching cost has an initial growth rate

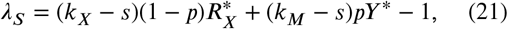

where -^*^ed variables indicate the steady state values before invasion (see SI 8 for more detailed analyses). If it is invading the original two-cooperator steady state (*X*^*^, *Y* ^*^), the condition for successful invasion is given by

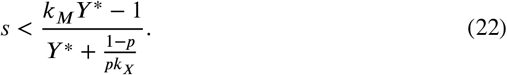

The dashed line in fig. 6A illustrates the contour of *λ* = 0 as *s* and *p* are varied. For values of *s* below this line, inequality (22) is satisfied and a cost-imposed mutant can invade. Mutants with higher *p* can tolerate higher switching costs *s*.

**Figure 6:**
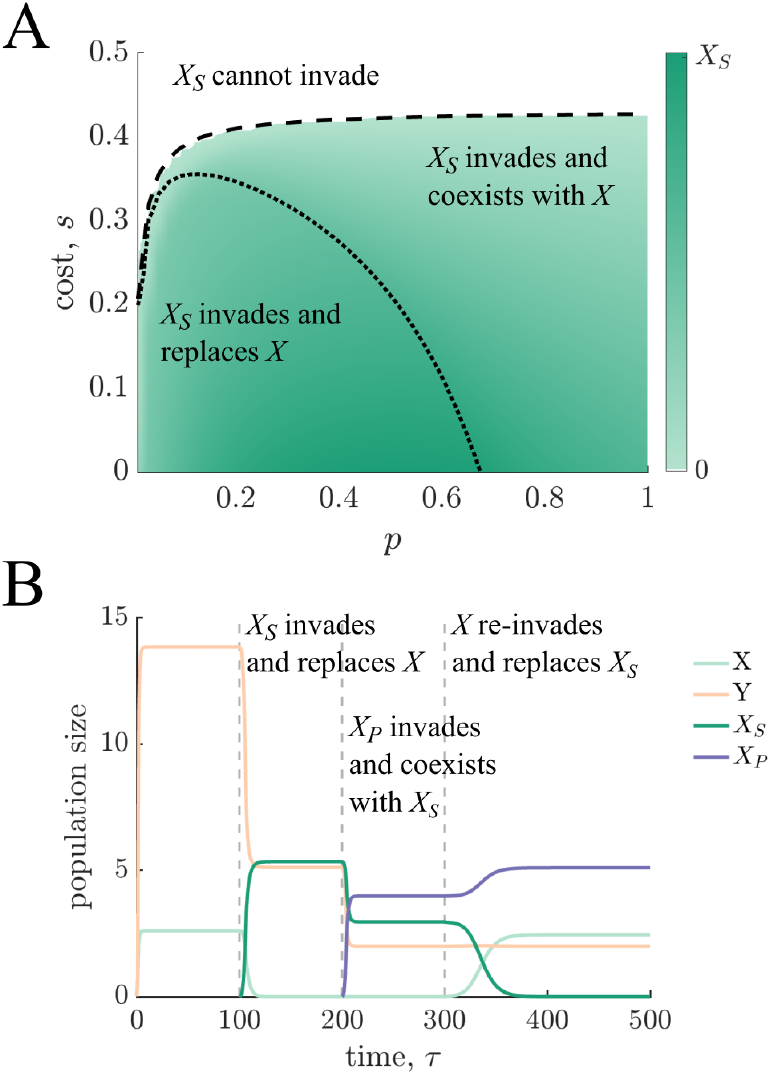
Invasion of a mixotrophic mutant with switching cost. A) Heat map of the steady state population of a switching cost-imposed mixotrophic mutant, *X*_*S*_, after invading the two- cooperator system with *X* and *Y* . Parameters *p* and *s* are varied and all others are as in fig. 2. The dashed line shows the invasion threshold where *λ* = 0: for higher costs and lower predation fractions the mutant cannot invade, while if the predation fraction is high enough given the cost, then the mutant can invade and establish in the community. The dotted line shows the critical threshold *p*_*crit*_(*s*) as a function of switching cost: when the predation fraction is above *p*_*crit*_(*s*) the mutant coexists with *X*, while below it outcompetes and replaces *X*. B) Plot shows an example where a cost-imposed mixotroph *X*_*S*_ invades the two-cooperator system with *X* and *Y* and replaces the original *X* species. Parameters are as in fig. 2, additionally *X*_*S*_ has predation fraction *p* = 0.3 and cost *s* = 0.16. At the new steady state a pure predator species *X*_*P*_ invades and creates a steady state which the original *X* species is able to re-invade. Once *X, Y*, and *X*_*P*_ have established a community, no cost-imposed (mixotroph) mutants can invade.

After *X*_*S*_ invades, the population dynamics are similar to those earlier when a cost-free mixotroph invades (see SI 9 and fig. 6A). One key difference in the cost-imposed system is that a pure predator can always invade a community with an *X*_*S*_ species. We outline all the cases in SI 9. Briefly, if *X*_*P*_ invades a steady state with *X, Y*, and *X*_*S*_, then the pure predator replaces the mixed strategy *X*_*S*_ mutant. Whereas, if *X*_*P*_ invades a steady state with *X*_*S*_ and *Y*, then all three (*Y*, *X*_*S*_, and *X*_*P*_) coexist. This new community structure, however, is not evolutionarily stable. The new equilibrium value of resource *R*_*X*_, makes the system susceptible to reinvasion by *X* (see SI 9). A re-invading *X* species replaces the *X*_*S*_ population, illustrated, for example, in fig. 6B. Thus, we find that it is not possible for *X, Y*, *X*_*S*_, and *X*_*P*_ to all coexist. Furthermore, once a community consists of *X, Y*, and *X*_*P*_ it cannot be invaded by any mixotroph *X*_*S*_. Hence, when mixotroph mutants have costs associated with switching between feeding strategies, the community evolves towards specialist species with single resources (*X, Y*, *X*_*P*_).

## 4. Discussion

A distinguishing feature of deeply-rooted unicellular eukaryotic cells is their ability to consume other cells through engulfment (Tikhonenkov et al., 2022; Al Jewari and Baldauf, 2023; Lamza, 2026). By contrast, very few prokaryotes are known that have phagocytic-like behaviour (Lage et al., 2025). Indeed, fewer prokaryote species probably engage in predation compared to the spectrum of other ecological interactions ranging from competition to cooperation (Vasse and Velicer, 2026). This suggests that in the evolutionary transition from prokaryotes to eukaryotes, phagocytic predation emerged from communities with primarily competitive or cooperative interactions. There are clear benefits to developing the ability to eat a competitor. However, another transition in eukaryogenesis was the evolution of endosymbiosis with an endosymbiont cell residing inside and coexisting with the host cell that engulfed it. In this context a mutual benefit between the host and endosymbiont cell seems more conducive to establishing a favourable endosymbiosis, raising the question of when it is beneficial to consume a cooperator. Here, we modelled the synergistic interaction between two species as a dynamical system and used invasion analysis to determine the conditions for a phagocytic mutant to invade. Fundamentally, we found the evolution of phagocytosis rested on the invading mutant receiving a sufficient growth benefit from predation, relative to the abundance of the prey population.

If our model so readily predicts the evolution of phagocytosis, even from a cooperative system, why then is predation by prokaryotes so rarely observed? One reason could be physical constraints in the machinery required. In order to phagocytose another species a cell has to be able to bind the prey to its surface, manipulate its body to engulf the prey, and then kill and digest the prey it has internalised (Cavalier-Smith, 2009). These adaptations all have associated energetic costs and it is not clear that they individually provide fitness benefits to the cell (Richards and Endres, 2017; Schavemaker et al., 2025). While there have been no observations of archaea that are capable of phagocytosis, genome sequences have identified some species with some of the necessary structural features (Panagiotou et al., 2026; Tobiasson et al., 2026). This has lead to popular theories that the eukaryotic cell stems from an archaeon with primitive phagocytic capabilities (Asgard archaea are a commonly cited candidate) that merged with the Alphaproteobacterium ancestor of mitochondria (Martijn and Ettema, 2013; Yutin et al., 2009; Mills, 2020). However, recent discoveries of bacteria (*Candidatus* Uabimicrobium amorphum and *Candidatus* Uabimicrobium helgolandensi) with phagocytic-like predation keep the door wide open to alternative origins for eukaryotic cells (Shiratori et al., 2019; Wurzbacher et al., 2024).

The physical requirements of phagocytosis may explain why its evolution from cells lacking this machinery is seemingly so rare. This has lead some to hypothesise that, during the transition from prokaryotes to eukaryotes, phagocytosis was preceded by endobiotic predation or parasitism (Davidov and Jurkevitch, 2009; Gassler et al., 2025). Under this hypothesis, a predatory prokaryote would enter its prey or host to consume and/or grow inside. Persistence of this predator could eventually lead to endosymbiosis; the development of this compartmental cell structure might lead to the evolution of machinery for the host cell to engulf or exchange endosymbionts. Our model captures phagocytosis primarily through the privatisation of resources by predatory mutants, and could therefore also be viewed as a highly simplified model of endobiotic predation. A more explicit representation of whether predators are inside or outside their prey would, however, be needed to capture this process more fully (such as, for example, (Summers and Kreft, 2022)). Further model modifications could be used to capture other forms of microbial predation and their evolution. For example, predation including external digestion would generate public goods. Such forms of predation might naturally emerge from competitive arms races incentivising the evolution of bacterial warfare over some limiting environmental factors. Given the prevalence of bacterial warfare mechanisms, some have suggested that such forms of bacterial predation might be much more common than we expect and their apparent rarity is merely a consequence of a lack of looking (Vasse and Velicer, 2026).

Within the context of phagocytosis and this model format, we found predatory mutants could both invade and coexist with the original cooperative community. Imposing a cost to the switching behaviour of mixotrophs which could both engage in predation and eat the same resource as their ancestor, eventually led to an evolutionarily stable community with three specialist species: the original two cooperators and a pure predator. Once the stable community established, no mixotrophs could invade. Even without the switching cost, there was a niche separation into two X-type species with a higher and lower predation fraction (above and below *p*_*crit*_, see fig. 5A). This suggests that the evolution of this system favours a separation into distinct trophic levels where each species is a specialist for different resources. While our model is a simplification of the evolutionary history leading to phagocytosis, this result provides another possible explanation for the rarity of phagocytosis in extant prokaryotes. Once a specialised predator has invaded and become established in the original cooperative community, our model predicts that selection no longer favours the invasion of mixotrophs, thus separating the evolutionary trajectories of species into distinct ecological niches.

In reality, microbes live in diverse communities where they are faced with multiple social and environmental pressures. Moreover, both these biotic and abiotic factors are intrinsically interconnected, as microbes alter their behaviour in response to each other and to their environment, and collectively this modifies that environment. This plasticity could lead to evolutionary trajectories beyond the specialist community structure predicted by our model, for example, creating opportunities for the invasion of cheaters, introducing fatal instabilities if an obligate partner becomes prey, or favouring bet-hedging strategies. Nevertheless, our model highlights that predation does not have to be predicated on an antagonistic interaction. Instead, we showed that predation can evolve from a cooperative community and the resulting predator can stably coexist within that community. We specifically modelled a syntrophic interaction, analogous to those found in microbial communities of deep-sea vents where primary degraders support the growth of secondary consumers (commonly methanogenic archaea) responsible for the efficient removal of waste products (Morris et al., 2013). These and other beneficial interactions are widespread across microbial communities, alongside a spectrum of antagonistic interactions (Foster and Bell, 2012; Granato et al., 2019; Kost et al., 2023; Morris et al., 2013), and which of these interactions arise can be highly context dependent (Di Martino et al., 2024; Solowiej-Wedderburn et al., 2025; Vasse et al., 2024). Exploring how predators arise, coexist, and interact across this dynamic landscape can help us uncover the processes that shape the evolution and biodiversity of microbial communities.

## Acknowledgements

J.S-W. and E.L. thank the Swedish Research Council (Vetenskapsrådet) for providing the means to conduct this work. J.G.O. thanks the Santa Fe Institute for hosting our working group meetings and providing office space and hospitality during a sabbatical visit in 2022. The authors would additionally like to thank Rebecka Thaysen for helpful discussions and IceLab for providing a supportive and collaborative environment for interdisciplinary science.

## Supporting information

### SI 1 Time rescaling of ODEs

In the main text we introduced ODEs (1)–(6) to describe the dynamics of a system of two cooperators *X* and *Y* . Here, we rescale time by the outflux constant *ϕ* to rewrite the system in terms of a dimensionless time parameter *τ*:= *ϕt*:

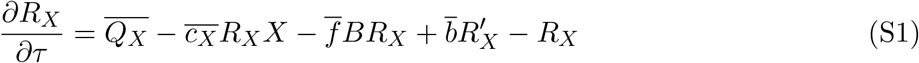

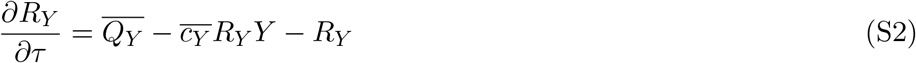

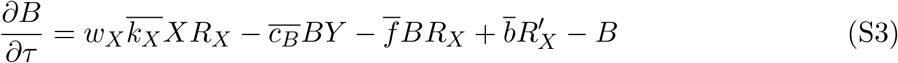

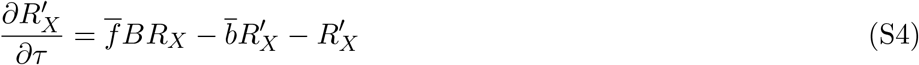

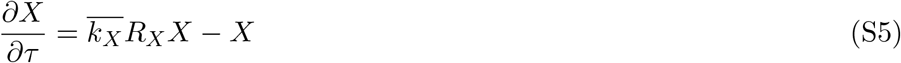

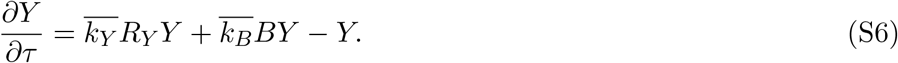

We have redefined all the remaining parameters according to this rescaling, for example, 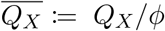. We then use this time rescaled system throughout the rest of the text but drop the bars for readability.

### SI 2 Steady states of the two-cooperator system

Here, we determine the steady states of the two-cooperator system of ODEs given by (S1)–(S6). Specifically, we are interested in the solutions that are biologically realistic with all resources and species ≥ 0. Setting equation (S5) = 0, we see that there is a solution with *X* = 0, similarly, (S6) = 0 is satisfied by *Y* = 0. We separately consider these steady states with *X* and/or *Y* = 0, and the coexistence steady state solution(s) with *X, Y* > 0.

#### Trivial steady state

The trivial steady state with *X* = 0 and *Y* = 0 is:

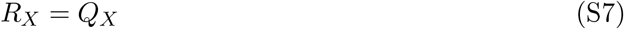

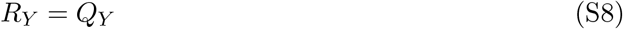

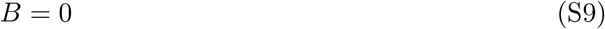

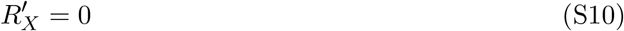

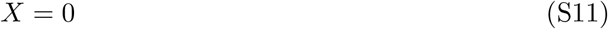

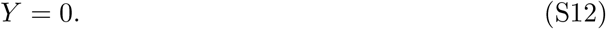

#### Existence of only one species

The steady state with *X* > 0 and *Y* = 0:

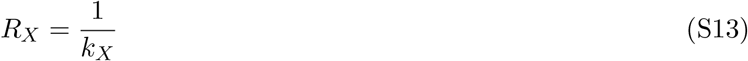

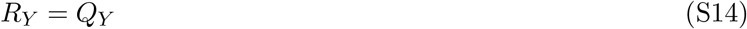

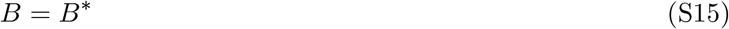

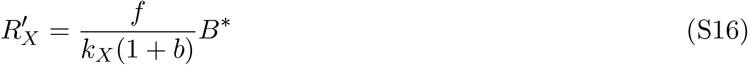

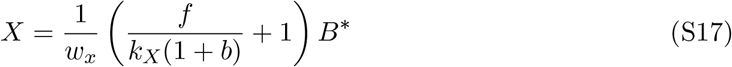

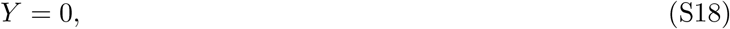

where the steady state value of inhibitor *B* is given by the expression

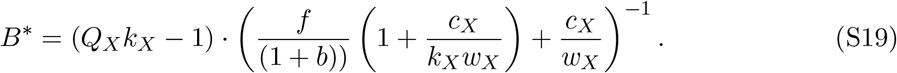

In order that *B*^∗^ is realistic, we require there to be enough influx of resource *R*_*X*_, i.e. *Q*_*X*_*k*_*X*_ > 1. If *B*^∗^ > 0 (i.e. is physical), we can immediately see that all other variables are also physical for any choice of positive parameters.

The steady state with *X* = 0 and *Y* > 0:

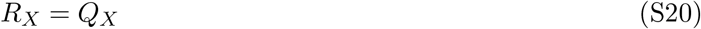

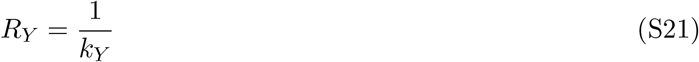

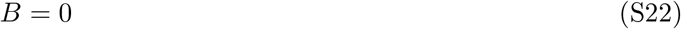

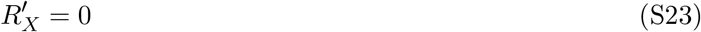

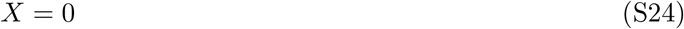

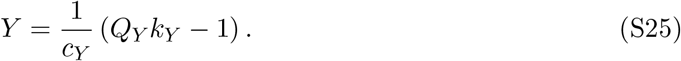

Hence, in the absence of *X* the only requirement for *Y* to survive is that there is a sufficient influx of resource *R*_*Y*_, i.e. *Q*_*Y*_ *k*_*Y*_ > 1.

#### Coexistence steady state

The coexistence steady state, where *X, Y* > 0, is given by:

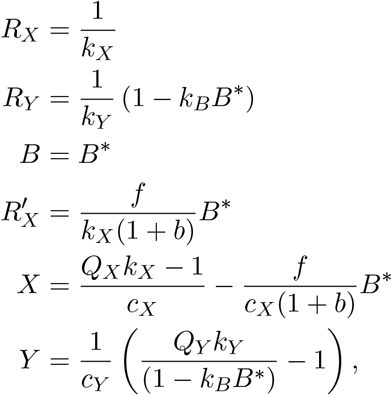

where *B*^∗^ is a solution to quadratic equation

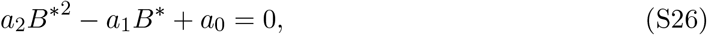

with

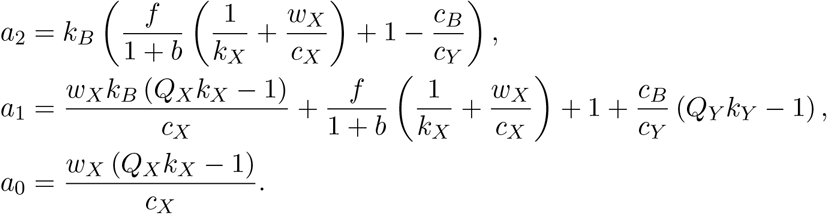

The quadratic equation (S26) for *B*^∗^ has real solutions as long as

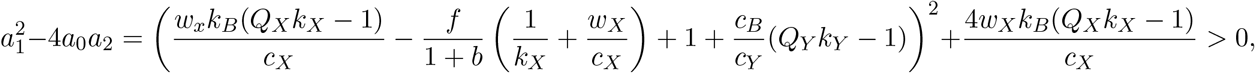

which is satisfied if *Q*_*X*_*k*_*X*_ > 1 as we have assumed that all parameters are ≥ 0.

For the coexistence steady state to exist, we need all species to have positive values and resources be non-negative. This requires 0 < *B*^∗^ < 1*/k*_*B*_. To determine when these constraints are satisfied, we rewrite the quadratic coefficients in terms of *k*_*B*_ as

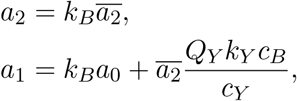

where 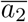 is a function of *f, b, k*_*X*_, *w*_*X*_, *c*_*X*_, *c*_*B*_, and *c*_*Y*_ . We note that 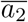 can be negative if *c*_*B*_ is sufficiently large relative to the other parameters.

Thus, *B*^∗^ is given by

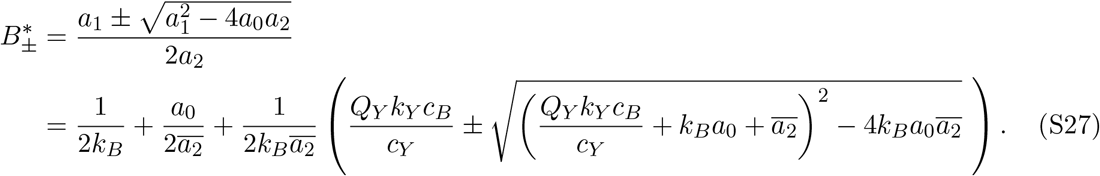

If 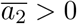, then we have

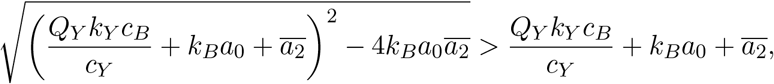

while if 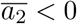, then we have

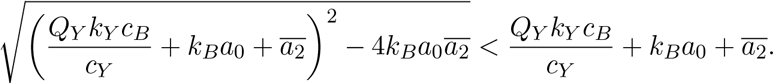

In both cases, combining these inequalities with expression (S27) for *B*^∗^ gives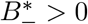. We can also rewrite the expression in the 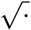 as

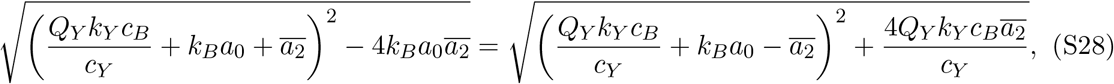

which satisfies 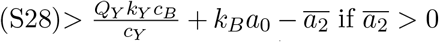 and 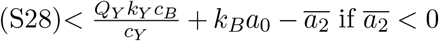. In both cases, these give that *B*−∗ < 1*/k*_*B*_.

Conversely, substituting expression (S28) into expression (S27) for *B*+∗, we find that if 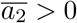,

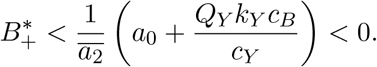

Alternatively, we can rewrite the expression in the 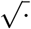 as

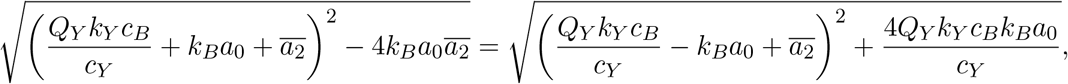

which, if 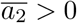, gives

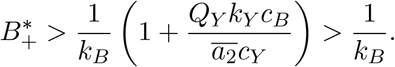

Hence, we have shown that 0 < *B*−∗ < 1*/k*_*B*_, while 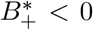 or 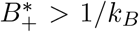 depending on the sign of 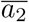 and so 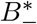 is the solution to (S26) that can give biologically realistic solutions. The solutions for *X* and *Y* give additional constraints on *Q*_*X*_ and *Q*_*Y*_ to ensure there are sufficient resources for the environment to support both species. Note that we need at least *Q*_*X*_*k*_*X*_ > 1.

#### Special case: *Y* **obligate on** *X*

There is a special case where there is no influx of resource *R*_*Y*_, i.e. *Q*_*Y*_ = 0.

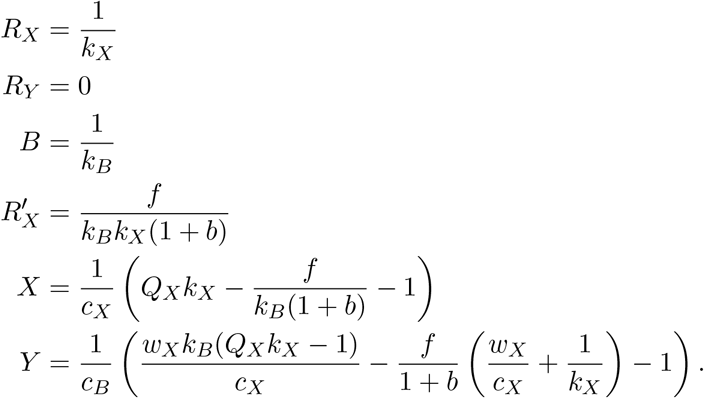

Thus, we see that if there is a sufficient influx of resource *R*_*X*_, i.e.

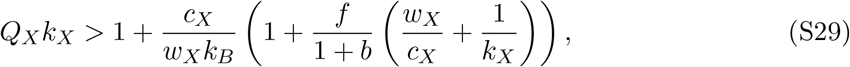

then we can have species *Y* surviving purely on the waste product produced by species *X*. This condition is also sufficient to ensure that the *X* population is positive. As the right-hand-side of inequality (S29) is greater than 1, the requirement for the influx of resource *R*_*X*_ to sustain both *X* and provide enough by-product for *Y* is greater than the requirement to just sustain *X*.

#### Stability of steady states

To determine the stability of the steady states we construct the Jacobian matrix for the system. For the two-cooperator system, this is given by

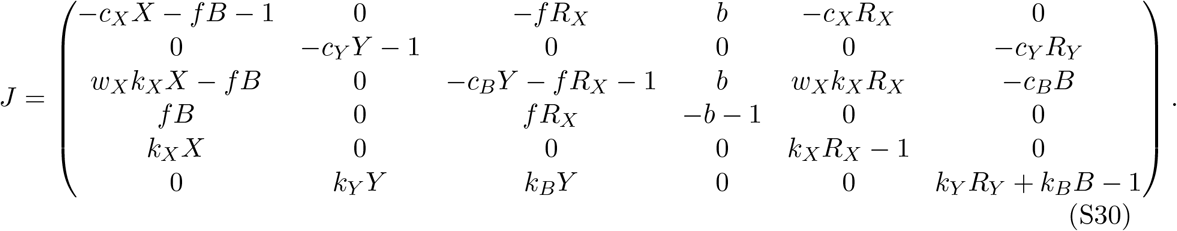

First we consider the fixed point with (*X, Y*) = (0, 0). Substituting expressions (S7)–(S12) for this steady state into (S30), we find that the eigenvalues of the Jacobian matrix satisfy

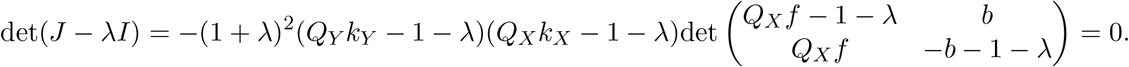

This means that the fixed point at (0, 0) has positive eigenvalues *λ*_1_ = *Q*_*Y*_ *k*_*Y*_ − 1 or *λ*_2_ = *Q*_*X*_*k*_*X*_ − 1 if the conditions are met for either fixed points with positive *X* or *Y* values. Hence, in these conditions the (0, 0) steady state is unstable.

Substituting expressions (S13)–(S18) for the (*X*^∗^, 0) > (0, 0) steady state into (S30), we find that the eigenvalues of the Jacobian matrix satisfy

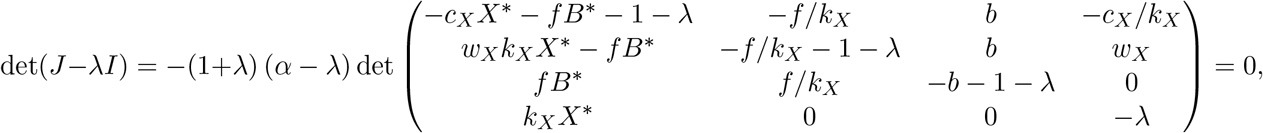

where *X*^∗^ and *B*^∗^ are given by (S17) and (S19), respectively, and

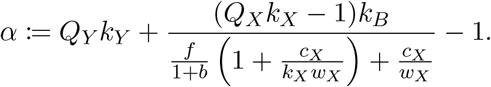

Thus, we see that two of the eigenvalues are *λ*_1_ = −1 > 0 and *λ*_2_ = *α*, where *α* > 0 if we are in a parameter regime where the (*X*^∗^, *Y* ^∗^) fixed point is > (0, 0). So in this parameter regime (*X*^∗^, 0) is a saddle.

Similarly, substituting expressions (S20)–(S25) for the (0, *Y* ^∗^) > (0, 0) steady state into (S30), we find that the eigenvalues of the Jacobian matrix satisfy

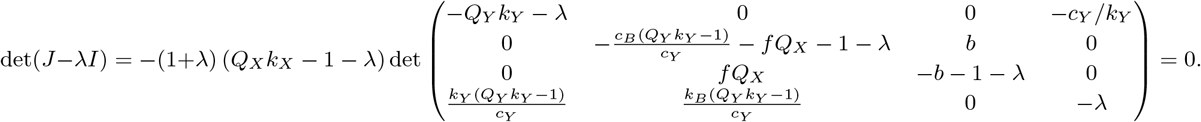

In this case, two of the eigenvalues are *λ*_1_ = −1 < 0 and *λ*_2_ = *Q*_*X*_*k*_*X*_ −1. Again, we find *λ*_2_ > 0 if we are in a parameter regime where (*X*^∗^, *Y* ^∗^) > (0, 0). Hence, if we have parameters such that the (*X*^∗^, *Y* ^∗^) fixed point is biologically realistic, we also find that both the steady states where only one can exist are unstable saddles. In these parameter regimes there is a sufficient influx of resources to push the system towards the steady state where (*X*^∗^, *Y* ^∗^) > (0, 0). Randomly sampling parameters in this regime we numerically confirm that the solutions converge to this coexistence steady state.

### SI 3 External influx of *B* for *X* obligate on *Y*

In SI SI 2 we determined the steady states of the two-cooperator system. There we considered the special case where *Y* has no resource influx (*Q*_*Y*_ = 0). In this case, it is impossible for *Y* to survive by itself, however, it could be possible for *Y* to grow in the presence of *X* forming an obligate interaction. (This requires a sufficient influx of *R*_*X*_ to support both *X* and *Y* populations.) We also showed that the condition for *X* to survive alone, *Q*_*X*_*k*_*X*_ > 1, is a necessary condition for *X* and *Y* to coexist. This means that there is no choice of parameters for which the coexistence steady state (*X, Y*) > (0, 0), but (*X*, 0) < (0, 0). In other words, we cannot have *X* obligate on *Y* under the dynamical system given by ODEs (1)–(6).

If we want to consider ecologies where *X* is obligate on *Y*, we could expand the system (1)–(6) to include an external influx of inhibitory compound *B*. We do this by rewriting equation (3) as

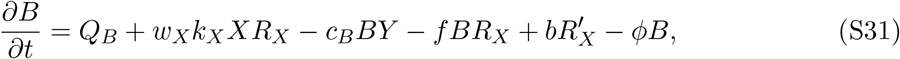

where *Q*_*B*_ is the influx of *B*. If *Q*_*B*_ = 0 then we retrieve the original system. We perform the same rescaling in time as in SI SI 1 to rewrite ODE (S31) as

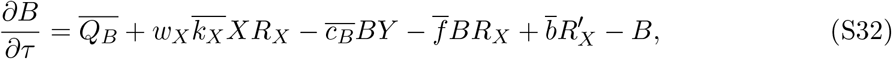

where again bars denote time-rescaled parameters, for example, *Q*_*B*_:= *Q*_*B*_*/ϕ*. We now use this time-rescaled system and drop the bars for readability.

To determine when *X* is obligate in this more general system, we first solve for the existence of *X* in the absence of *Y* . Here, the steady state is given by

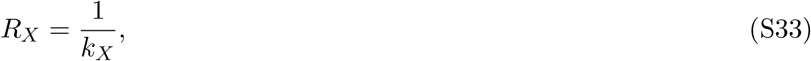

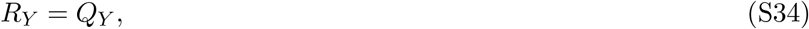

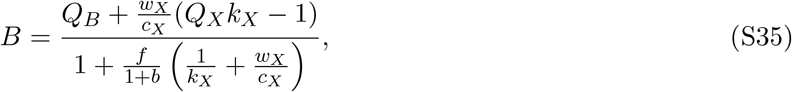

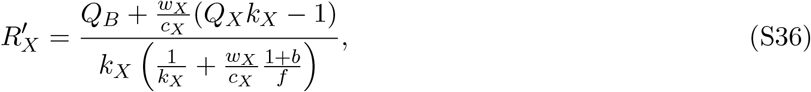

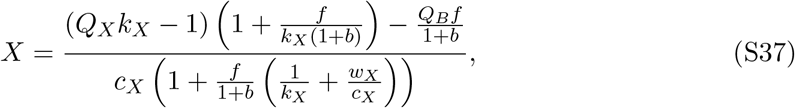

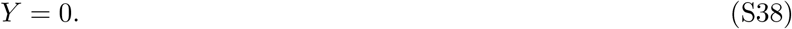

Hence, we see that if

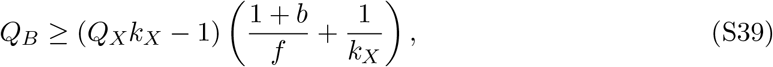

then expression (S37) for *X* is ≤ 0 and so *X* cannot survive alone.

Second, we solve for the coexistence steady state:

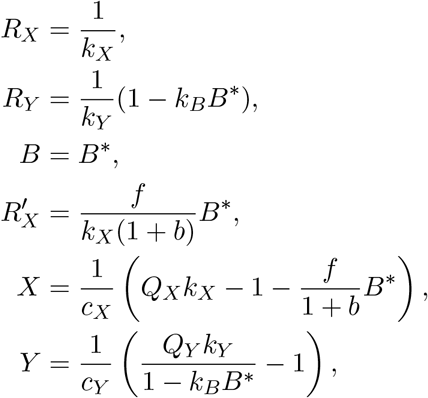

where *B*^∗^ is a solution to the quadratic equation where *B*^∗^ is a solution to quadratic equation

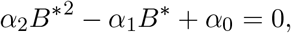

with

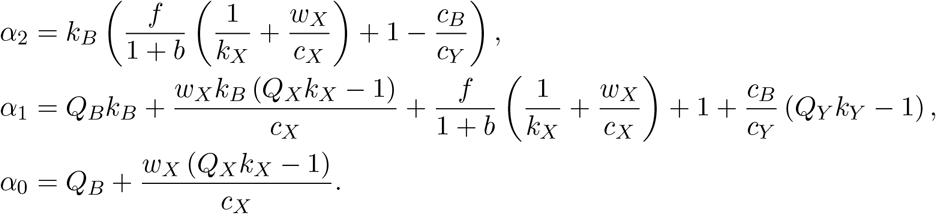

In parameter regimes where the coexistence steady state has positive solutions and *Q*_*B*_ satisfies inequality (S39) it is only possible for *X* to grow in the presence of *Y* . We illustrate an example case in fig. S1.

**Figure S1:**
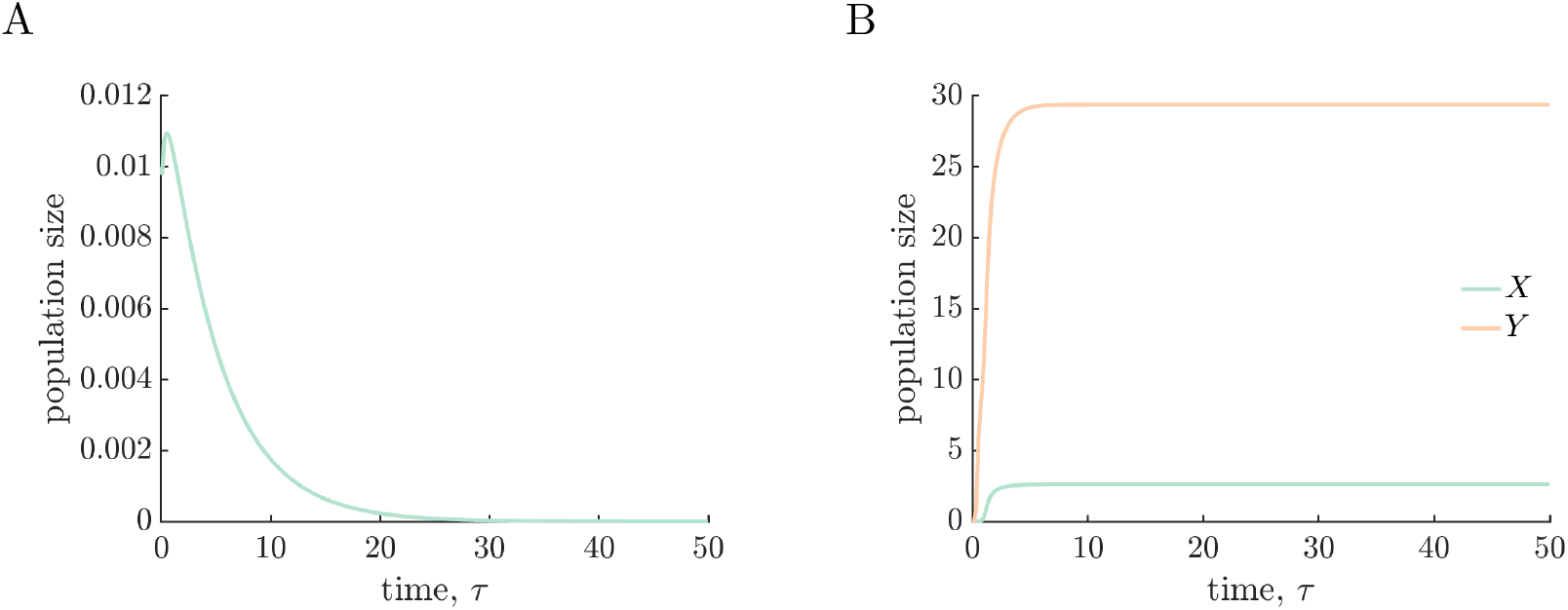
*X* obligate on *Y* . A) Plot of *X* population size alone with an external influx of *B*. Here *Q*_*B*_ = 30, and other parameters as as in fig. 2 in the main text. *X* cannot grow alone. B) Plot of *X* population in the same conditions but in the presence of *Y* species. *X* and *Y* can grow together, so *X* in these conditions is obligate on *Y* .

### SI 4 Steady state with a mutant

Now we consider the steady states of the system of ODEs given by (7)–(13) in the main text for two cooperators (*X* and *Y*) with a mutant species (*X*_*M*_). If the mutant fails to invade *X*_*M*_ = 0 and we have the steady states for the two cooperator system (see SI SI 2).

The steady state with coexistence of all three species (*X, Y, X*_*M*_) > (0, 0, 0) is given by:

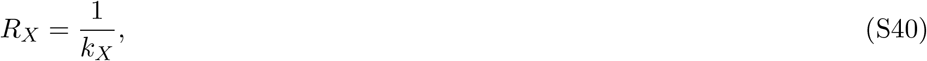

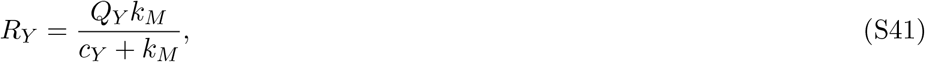

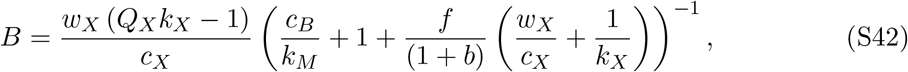

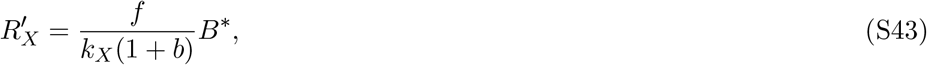

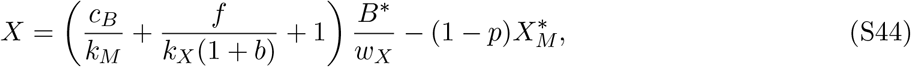

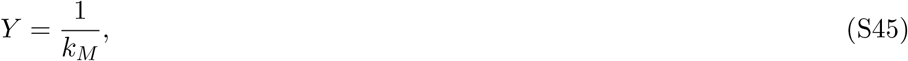

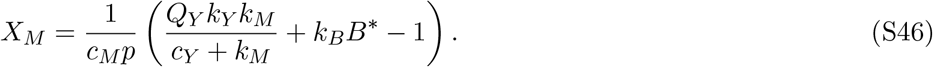

where *B*^∗^:= *B* and 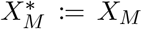 at this steady state. Biologically realistic solutions depend on enough resources and *p* sufficiently large that *X* and *X*_*M*_ can coexist. Selecting parameters such that *Q*_*X*_*k*_*X*_ > 1 ensures *B*^∗^ > 0 and 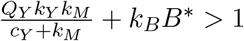 ensures *X*_*M*_ > 0. For *p*, we can determine a critical

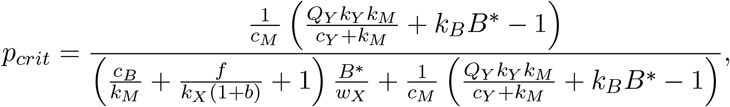

such that, if the resource requirements are met and *p* > *p*_*crit*_, *X* > 0 (expression (16) in the main text). In these cases, we have the coexistence of all three species *X, X*_*M*_, and *Y* at steady state. Otherwise, if *p* < *p*_*crit*_, a steady state with the coexistence of all three species does not exist and we find that *X*_*M*_ outcompetes *X*. It is possible to find analytic expressions for the coexistence of only *X*_*M*_ with *Y* by setting *X* = 0 in ODEs (7)–(13) and solving for the steady state.

Note that in expression (16) for *p*_*crit*_, the numerator is smaller than denominator (as long as *B*^∗^ > 0), so *p*_*crit*_ < 1; i.e. we don’t have situations where the condition for coexistence is with a predation probability ≥ 1. Furthermore, *p* = 1 is always greater than *p*_*crit*_, hence, as long as the conditions are met for invasion, a pure predator *X*_*M*_ can always coexist with *X* and *Y* .

We additionally note that the population of *Y* is independent of *p* and depends only on *k*_*M*_ (the growth benefit *X*_*M*_ receives from consuming *Y*); a higher *k*_*M*_ corresponds to a smaller *Y* population and larger *X*_*M*_ population. The requirement for the invasion of *X*_*M*_, setting *λ* > 0 in expression (15), ensures that *Y*_post_ = 1*/k*_*M*_ < *Y* ^∗^, where *Y* ^∗^ is the two-cooperator steady state population.

### SI 5 Evolutionary response of *Y*

In the main text, we consider the evolutionary response of *Y* via adaptation in the growth rate, 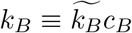, when consuming by-product *B*. We do this by testing the invasion of mutant *Y*_*M*_ with growth rate 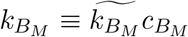 . When *Y*_*M*_ invades it replaces *Y* ; it is not possible to find a steady state solution where the two species coexist (i.e., there is no solution where *Y* ^∗^ > 0 and 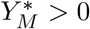. So the new steady states following a successful invasion by *Y*_*M*_ can be determined as in the previous system but replacing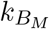 (and 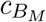) with the new 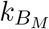 (and 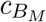) parameter(s). There are a few different outcomes for the community dynamics following the successful invasion of mutant *Y*_*M*_ depending on the steady state that it is invading and the underlying parameters; we summarised them in the main text and here outline them more methodically.

If *Y*_*M*_ invades a community of *Y* and one *X*-type species (either *X* or the mixotrophic *X*_*M*_), it leads to an increase in the *X* (or *X*_*M*_) population (this can be seen by substituting 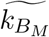 or 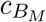 into the steady state expressions for *X* or *X*_*M*_ found in SI SI 2 and fig. S2A for *X* and fig. 4A for *X*_*M*_). If *Y*_*M*_ invades a community with all three species *X, X*_*M*_, and *Y*, then the outcome also depends on whether the adaptation in 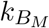 manifests as an adaptation in 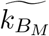 or 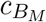 . An increase in 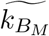 results in prey with a higher growth rate, which translates to an increase in the population size of *X*_*M*_ . This added benefit to *X*_*M*_ gives it a greater competitive advantage over *X* and so the *X* population decreases. The critical threshold *p*_*crit*_ is also dependent on *k*_*B*_; so when *Y*_*M*_ invades we define a new *p*_*crit, new*_ in terms of 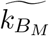 and 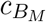 (see fig. S2B). If 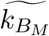 is large enough that *p*_*crit, new*_ > *p* > *p*_*crit*_, the invasion of *Y*_*M*_ gives *X*_*M*_ enough added benefit that it drives *X* extinct (see, e.g., fig. 4B and fig. S2B and C). There is an exception when *X*_*M*_ is a pure predator (*p* = 1); in this case it no longer competes for resources with *X* and the steady state *X* population is independent of 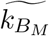 (see SI SI 4 and fig. S2D). Conversely, when *X*_*M*_ is a pure predator, an increase in 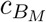 would correspond to an increase in both *X*_*M*_ and *X* populations (fig. S2D). Otherwise, when 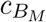 increases with a mixotrophic *X*_*M*_, there is a trade off between the increased removal of inhibitory by-product *B* and the competitive advantage *X*_*M*_ has with faster growing prey that means the *X* population could increase or decrease depending on the underlying parameters (fig. S2C).

Whether the resultant *Y*_*M*_ population steady state is greater or the same as *Y* ‘s population size prior to invasion depends on whether the parameters are such that *X* and *X*_*M*_ coexist post-invasion or not. In all the cases 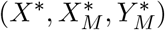 coexist, the 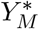 steady state population is independent of *k*_*B*_ (see SI SI 4), so 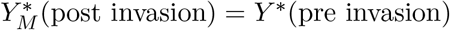. Whereas, if only *X*_*M*_ and *Y*_*M*_ coexist at steady state then 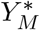 (post invasion) > *Y* ^∗^(pre invasion) (illustrated, e.g., in fig. 4A and B). Furthermore, *X*_*M*_ scales linearly with *k*_*B*_ so *X*_*M*_ always benefits from the invasion of *Y*_*M*_ (see SI SI 4). So if all three species can coexist, the evolutionary response of *Y* may result only in an increase in the *X*_*M*_ population size. This means that, while it is always possible for *Y*_*M*_ to invade with higher 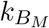, it only experiences the long term population size benefits if *X*_*M*_ ‘s predation fraction is small enough.

### SI 6 Steady state with two mutants

If *X*_*M*_1 invaded with *p*_1_ < *p*_*crit*_, then it would replace the original *X* population. This leaves a community that is capable of having two mutant species coexisting with the *Y* population. Here, we adapt the system of ODEs (7)–(13) to include a second mutant species and set the *X* population to 0. The resulting system is

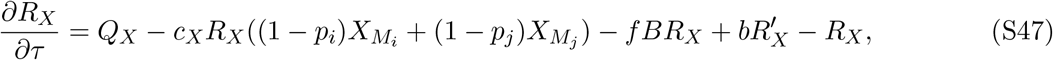

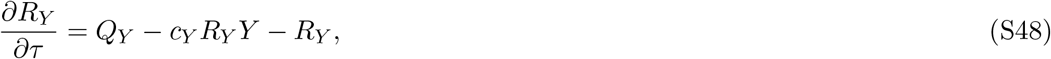

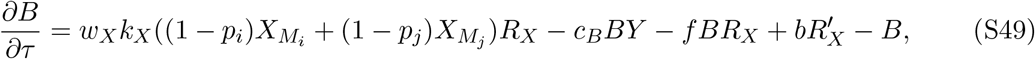

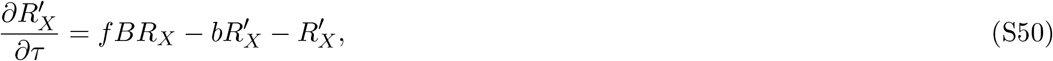

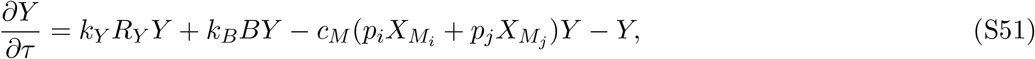

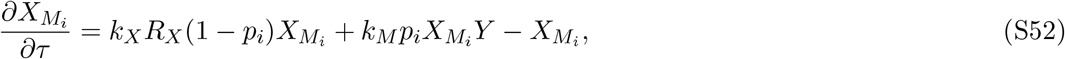

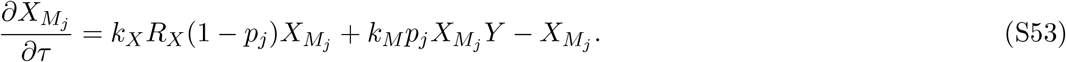

**Figure S2:**
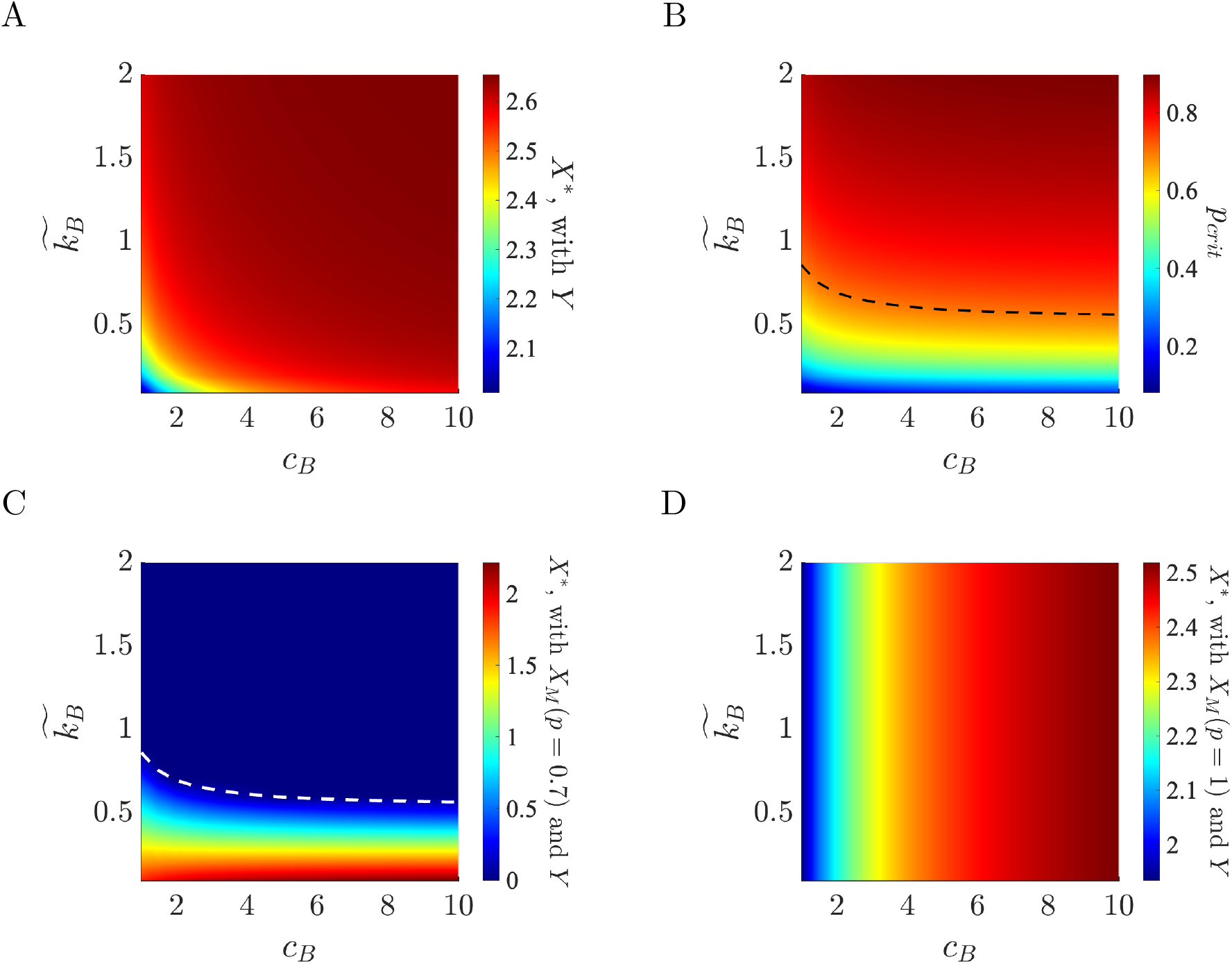
Effect of the evolutionary response of *Y* on *X*. A) Heatmap of the steady state *X* population with only *Y* species against 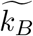 and *c*_*B*_. The *X* population is greater when either 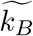 or *c*_*B*_ are larger, so if *Y* improves either of these mechanisms, this would lead to greater cooperative reward for *X*. B) Heatmap of the critical predation fraction *p*_*crit*_ against 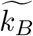 and *c*_*B*_. This shows that *p*_*crit*_ is greater for larger 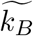 or *c*_*B*_, so if either of these parameters are increased, then there is a range, such that if *X*_*M*_ (*p* ∈ [p_*crit*_, *p*_*crit,new*_]), then the invasion of *Y*_*M*_ will drive the existing *X* species extinct. C and D) Heatmaps of the steady state *X* population with both *Y* and *X*_*M*_ species against 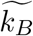 and *c*_*B*_. In (C) *X*_*M*_ has *p* = 0.7; for parameters 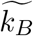 and *c*_*B*_ that give *p*_*crit*_ > 0.7, *X*^∗^ = 0. The effect of changing *c*_*B*_ on *X*^∗^ depend on 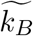. An increase in 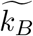 corresponds to a decrease in *X*^∗^ = 0, so if *Y*_*M*_ invades with higher 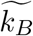 this would result in a decrease in *X* population at the new steady state. In (D) *X*_*M*_ has *p* = 1; in this case, *X*^∗^ is independent of 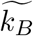, while increasing corresponds to an increase in *X*^∗^. So when coexisting with a pure predator, if adapts its growth conversion ratio 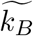, this would have no long term effect on *X* population (and new 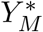 would be the same as *Y* ^∗^), whereas adapting the feeding rate *c*_*B*_ would at least result in an increase in *X*^∗^. In all panels, any other parameters are as in fig. 2 in the main text.

We calculate the coexistence steady state with *Y*, *X*_*M*_*i* and *X*_*M*_*j*, by setting all the ODEs (S47)–(S53) to zero and find

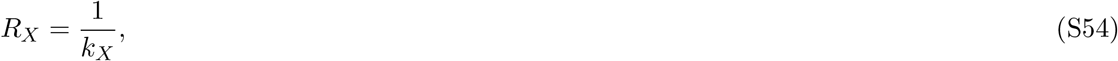

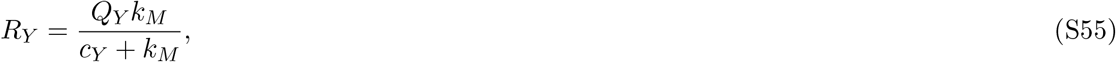

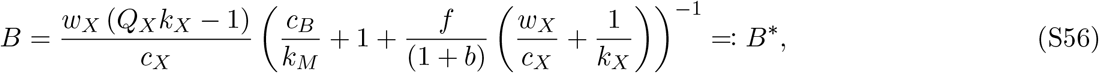

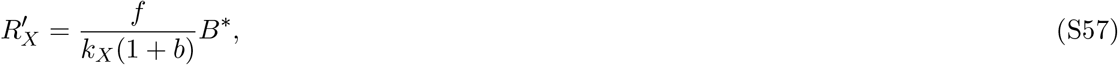

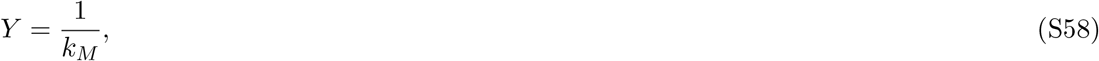

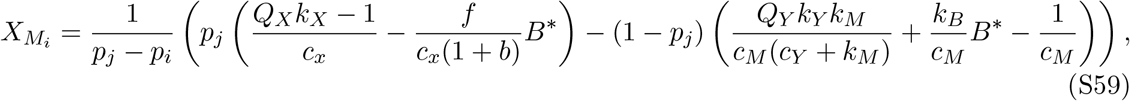

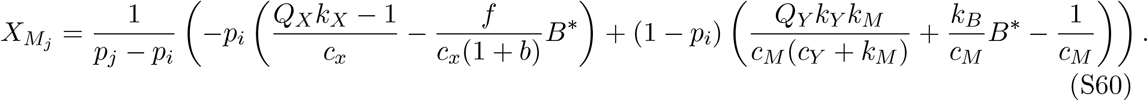

### SI 7 Degeneracy with multiple mutants

All the previous community models have been a subset of the general system of ODEs

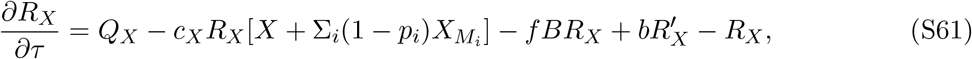

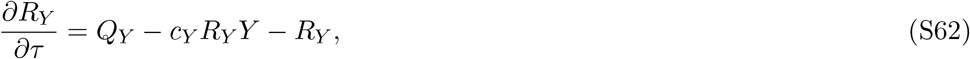

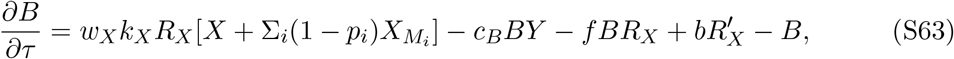

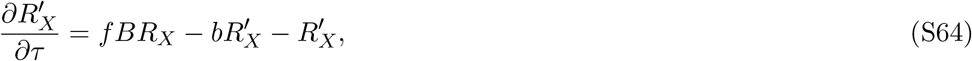

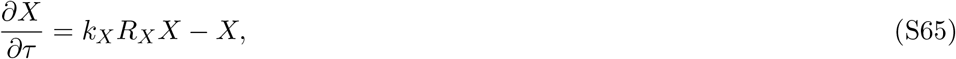

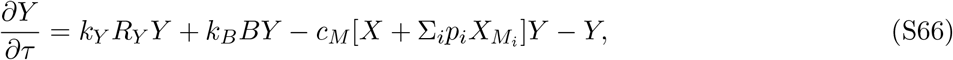

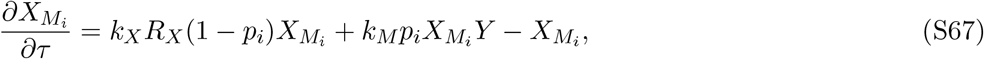

where *i* ∈ 1, 2, …, *N* for a community with *N* mutant species with different predation fractions *p*_*i*_. Solving this general system for the coexistence steady state solution of all species, we find that the resources are all given by the same expressions as when we have a system with three coexisting species (either *X, Y*, and *X*_*M*_ as in (S40)–(S43), or *Y*, 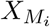, and 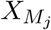 as in (S54)– (S57)). The *Y* population is also given by 1*/k*_*M*_ at steady state. However, there is a degeneracy in the system for *X* and *N* mutant 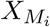 species. As such, at steady state we find that there are infinite solutions for *X* and 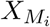 with their combined populations given by the hyperplane

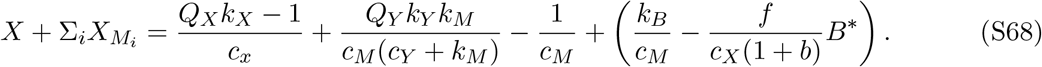

### SI 8 Invasion by mutant with switching cost

#### System of ODEs when mixotrophs have a switching cost

We modify the system of ODEs (7)–(13) to impose a switching cost, *s*, on the growth rate of mutants that can both consume resource *R*_*X*_ and predate on *Y* . This changes the equations of both the mixotrophs who experience the cost and and *B* which is produced by the mixotrophs at a rate proportional to their growth on *R*_*X*_. Additionally, we introduce new notation to distinguish mixotrophs with a switching cost (*X*_*S*_) from their cost-free counterparts in the previous model (*X*_*M*_). Whereas previously pure predators could be considered a special case of mutants with *p* = 1, now we consider pure predators separately as we assume that their singular feeding strategy does not incur this switching cost, and so denote them *X*_*P*_ . Together, we write the full system of ODEs when mixotrophs have a switching cost as

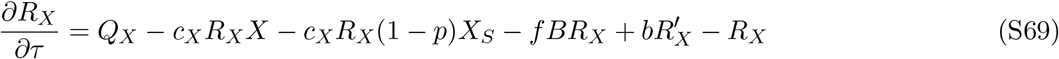

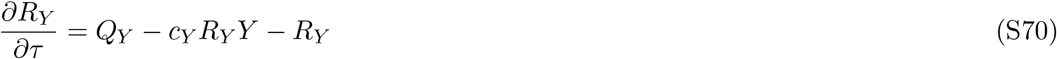

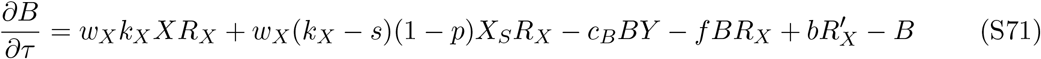

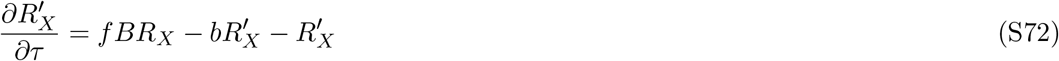

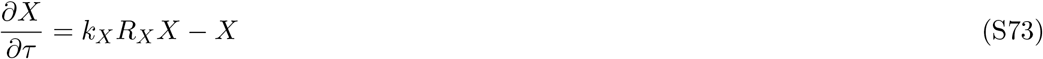

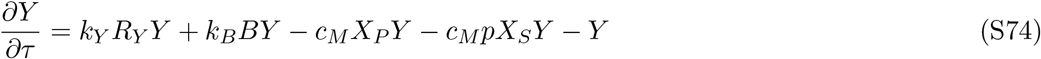

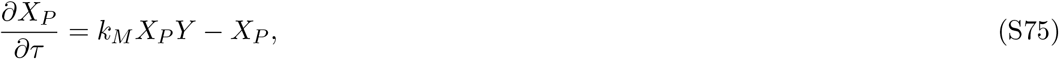

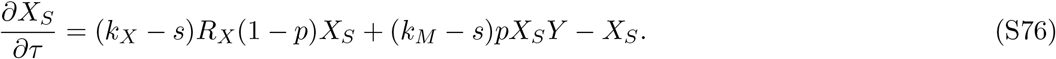

#### Conditions for invasion of the two-cooperator system

Under the assumptions of adaptive dynamics, we gave the initial growth rate of an invading mutant *X*_*S*_ with a switching cost as expression (21) in the main text. If it is invading the original two-cooperator steady state (*X*^∗^, *Y* ^∗^), the condition for successful invasion is given by

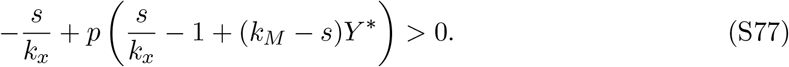

If *p* = 0, then the inequality (S77) cannot be satisfied for any positive switching cost *s*, as *X*_*S*_ will always be outcompeted by *X*. While if *p* ∈ (0, 1), we can rearrange the inequality (S77) to determine the upper bound for the maximum switching cost such that the mutant can invade given in the main text by (22). For a successful invasion of *X*_*S*_, we also require that it has *k*_*M*_ > 1*/Y* ^∗^, as in the case of an *X*_*M*_ mutant invasion (15).

### SI 9 Steady states following invasion by mutant with switching cost

#### Steady state with *X, Y*, and *X*_*S*_

First we consider the coexistence steady state after *X*_*S*_ invades the two-cooperator system. Setting ODEs (S69)–(S74) and (S76) to 0, we solve to find the coexistence steady state as

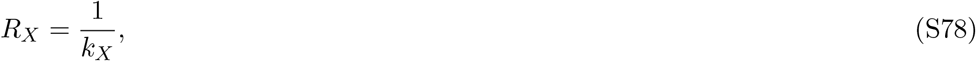

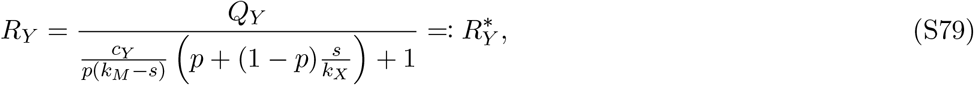

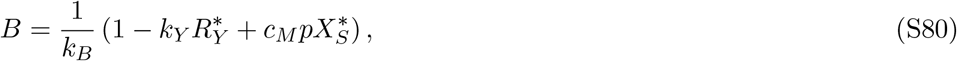

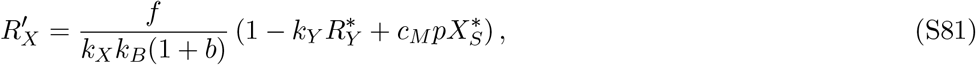

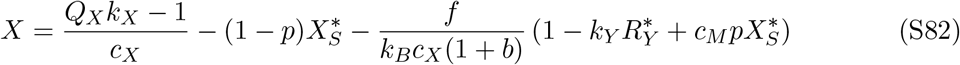

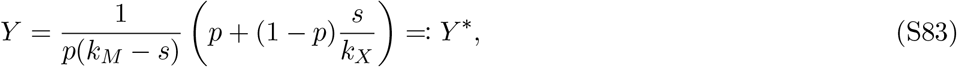

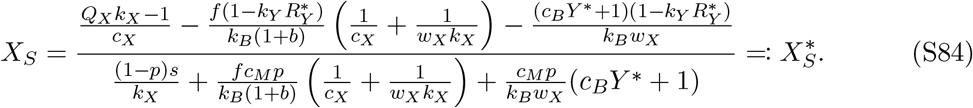

In order to realise the coexistence steady state, the values of *p* and *s* must be such that all the resource and population values given by expressions (S78)–(S84) are positive. We illustrate this for one set of parameter values with the dotted line in fig. 6A in the main text. For *p* and *s* values above this line *X*_*S*_ coexists with *X* and *Y* if it can invade. Whereas, for values below this line invading *X*_*S*_ population replaces *X*.

#### Invasion of subsequent 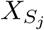 mutants

Subsequent 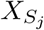 mutants can invade if their initial growth rate satisfies 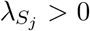, where 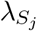 is given by expression (21) with *p* = *p*_*j*_. This depends on the steady state values of *R*_*X*_ and *Y* prior to invasion. If *p*_*i*_ > *p*_*crit*_(*s*), then we have the coexistence steady state prior to invasion, so *R*_*X*_ and *Y* are given by expressions (S78) and (S83), respectively. Substituting these expressions into (21) we find

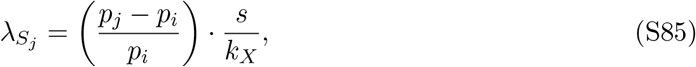

so 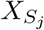 can invade if *p*_*j*_ > *p*_*i*_. However, if we try to solve for the steady state of ODEs (S69)– (S74) with an additional ODE for 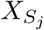 (identical to (S74) with *p* = *p*_*j*_), we find that it is not possible to have a steady state where *X*, 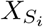, and 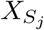 coexist. We can see this as setting equation (S73) to 0 with *X ≠* 0 implies that *R*_*X*_ = 1*/k*_*X*_, while doing the same with equation (19) for 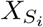 and 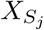 implies that *R*_*X*_ = 1*/*(*k*_*X*_ − *s*) for both 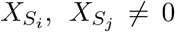, contradicting *R*_*X*_ = 1*/k*_*X*_. Instead numerical simulations show that if 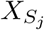 invades, then it replaces the 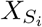 population, as illustrated, for example, in fig. 6B.

Alternatively, if *p*_*i*_ < *p*_*crit*_(*s*), then new mutant 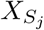 is invading a steady state with only *Y* and 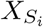 species. In this case, the steady state equation for 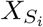(19) gives

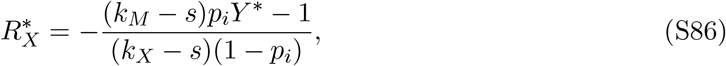

where *Y* ^∗^ is the steady state for species *Y* in this scenario. Substituting expression (S86) for 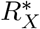 into expression (S85) for the initial growth rate of 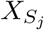, we find

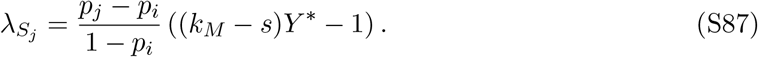

Again we see that a further mutant can invade if *p*_*j*_ > *p*_*i*_. We also note that expression (S87) is the same as expression (18) in the main text for the initial growth rate of a second cost-free mutant, but with *k*_*M*_ replaced by (*k*_*M*_ − s). In line with this, we find the invasion of subsequent mutants to follow similar dynamics in this case with 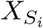 and 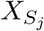 saturating the ecological niche so that the system becomes evolutionarily neutral to invasion by further 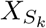 species.

#### Invasion by a pure predator *X*_*P*_

A pure predator can invade if *λ*_*P*_:= *k*_*M*_ *Y* ^∗^ −1 > 0, where *Y* ^∗^ again represents the steady state *Y* population prior to invasion. As previously, the value of *Y* ^∗^ depends on which steady state *X*_*P*_ is trying to invade. In the main text we showed that *X*_*P*_ can invade the two-cooperator system provided 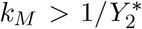, where we specify 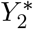 as the coexistence steady state of *X* and *Y* with no mutants.

If a cost-imposed mutant has invaded and can coexist with *X* and *Y*, then *Y* ^∗^ is given by expression (S83). This gives

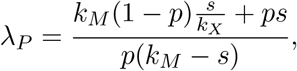

which is strictly positive as long as the conditions are satisfied for this steady state to exist (namely, (*k*_*M*_ − *s*) > 0 is a necessary condition for the coexistence of *X, Y*, and *X*_*S*_). Once *X*_*P*_ successfully invades, we set the full system of ODEs (S69)–(S76) to 0 to determine whether all four species can coexist at steady state. If *X, Y, X*_*S*_, *X*_*P*_ *≠* 0 then (S73) gives *R*_*X*_ = 1*/k*_*X*_ and (20) gives *Y* = 1*/k*_*M*_ . Substituting these into expression (19), we obtain (*k*_*M*_ (1 − *p*) + *pk*_*X*_)*s/k*_*X*_*k*_*M*_ which is strictly positive for *p* ∈ (0, 1) and *s, k*_*M*_, *k*_*X*_ > 0, contradicting the steady state condition that this expression should be 0. Hence, we see that *X, Y*, *X*_*P*_, and *X*_*S*_ cannot all coexist at steady state. Instead, we find that *X*_*P*_ replaces *X*_*S*_.

Finally, *X*_*P*_ could invade a steady state with *Y* and mutant(s) *X*_*S*_*i* (but *X* = 0). In analogy to the steady state with multiple cost-free mutants *X*_*M*_*i*, the steady state *Y* population with mutant(s) *X*_*S*_*i* is *Y* ^∗^ ≥ 1*/*(*k*_*M*_ −*s*), where we again require *k*_*M*_ > *s*. This gives *λ*_*P*_ ≥ *s/*(*k*_*M*_ −*s*) > 0. Hence, in all cases a pure predator can invade.

#### Coexistence of *Y*, *X*_*S*_, and *X*_*P*_

Just as it is not possible for *X, Y*, *X*_*P*_, and *X*_*S*_ to all coexist, it is also not possible for multiple mutants 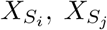, … to coexist with *Y* and *X*_*P*_ . We can see this as the steady state solution for multiple 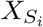 implies *Y* ^∗^ = 1*/*(*k*_*M*_ − *s*), in contradiction to setting ODE (20) for *X*_*P*_ to steady state which implies *Y* ^∗^ = 1*/k*_*M*_ . Hence, if *X*_*P*_ invades a system with multiple cost-imposed mutants it replaces their population(s). However, it is possible for *X*_*P*_ to coexist with one cost-imposed mutant population and *Y* . In this case, the steady state is given by

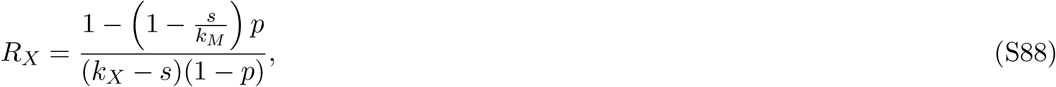

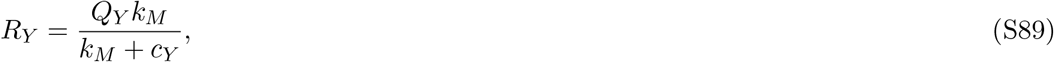

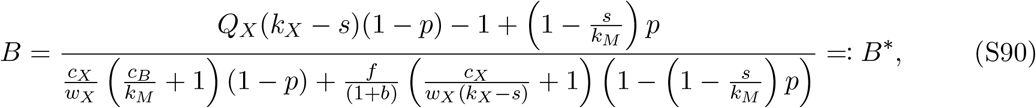

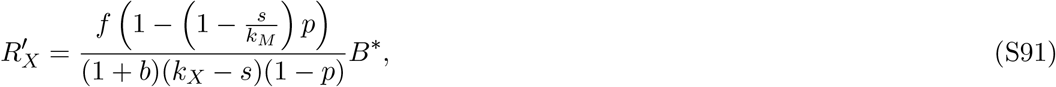

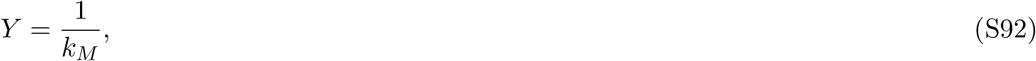

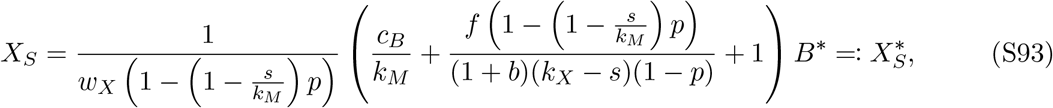

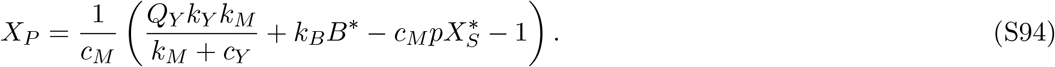

#### Reinvasion by *X*

We can see above that if a pure predator *X*_*P*_ invades a system with *Y* and cost-imposed mutant *X*_*S*_, this shifts the steady state concentration of resource *R*_*X*_. Consequently, it is possible for the original *X* species that was replaced by *X*_*S*_ to re-invade the system. We can see this using adaptive dynamics as the initial growth rate of *X* would be given by *λ*_*X*_ = *k*_*X*_*R*_*X*_ − 1. Substituting (S88) into this expression, we obtain *λ*_*X*_ = *s* (*k*_*X*_*p* + *k*_*M*_ (1 − *p*)) */k*_*M*_ (*k*_*X*_ − *s*)(1 − *p*) and as all these terms are positive we have *λ*_*X*_ > 0 so *X* is able to successfully re-invade a steady state with *X*_*S*_, *X*_*P*_, and *Y* .

#### System with *X, Y*, and *X*_*P*_ is robust to invasion by *X*_*S*_

The steady state with *X, Y*, and *X*_*P*_ is given by expressions (S40)–(S46), where the solution for *X*_*P*_ given by *X*_*M*_ with *p* = 1. To determine whether cost-imposed mutant *X*_*S*_ can invade this steady state, we substitute these expressions into the initial growth rate *λ*_*S*_ given by (21). This gives *λ*_*S*_ = −*s* ((1 − *p*)*/k*_*X*_ + *p/k*_*M*_) < 0, showing that it is not possible for a cost-imposed mutant to invade once *X, Y*, and *X*_*P*_ have established.

